# Genome analyses show strong selection on coloration, morphological and behavioral phenotypes in birds-of-paradise

**DOI:** 10.1101/287086

**Authors:** Stefan Prost, Ellie E. Armstrong, Johan Nylander, Gregg W.C. Thomas, Alexander Suh, Bent Petersen, Love Dalen, Brett W. Benz, Mozes P.K. Blom, Eleftheria Palkopoulou, Per G.P. Ericson, Martin Irestedt

**Author notes:** Corresponding authors: Stefan Prost, Martin Irestedt.

## Abstract

The diverse array of phenotypes and courtship displays exhibited by the birds-of-paradise have long fascinated scientists and laymen alike. Remarkably, almost nothing is known about the genomics of this iconic radiation. There are 41 species in 16 genera currently recognized within the birds-of-paradise family (*Paradisaeidae*), most of which are endemic to the island of New Guinea. In this study, we sequenced genomes of representatives from all five major clades within this family to characterize genomic changes that may have played an important role in the evolution of the group’s extensive phenotypic diversity. We found genes important for coloration, morphology and feather development to be under positive selection. GO enrichment of positively selected genes in the birds-of-paradise showed an enrichment for collagen, glycogen synthesis and regulation, eye development and other categories. In the core birds-of-paradise, we found GO categories for ‘startle response’ (response to predators) and ‘olfactory receptor activity’ to be enriched among the gene families expanding significantly faster compared to the other birds in our study. Furthermore, we found novel families of retrovirus-like retrotransposons active in all three *de novo* genomes since the early diversification of the birds-of-paradise group, which could have potentially played a role in the evolution of this fascinating group of birds.

## Background

> *‘Every ornithologist and birdwatcher has his favourite group of birds. Frankly, my own are the birds of paradise and bowerbirds. If they do not rank as high in world-wide popularity as they deserve it is only because so little is known about them.’*
>
> — *Ernst Mayer* (*in Gilliard 1969 [1]*)

The spectacular morphological and behavioral diversity found in birds-of-paradise (*Paradisaeidae*) form one of the most remarkable examples in the animal kingdom of traits that are thought to have evolved via forces of sexual selection and female choice. The family is comprised of 41 recognized species divided into 16 genera [2], all of which are confined to the Australo-Papuan realm. The birds-of-paradise have adapted to a wide variety of habitats ranging from tropical lowlands to high-altitude mountain forests [3] and in the process acquired a diverse set of morphological traits, some of which specifically fit their ecology and behavior. Some species are sexually monomorphic and crow-like in appearance with simple mating systems, whereas others have complex courtship behaviors and display strong sexual dimorphism with males exhibiting elaborate feather ornaments that serve as secondary sexual traits [3]. As such, strong sexual and natural selection have likely acted in concert to produce the exquisite phenotypic diversity among members the *Paradisaeidae*.

While having attracted substantial attention from systematists for centuries, the evolutionary processes and genomic mechanisms that have shaped these phenotypes remain largely unknown. In the past, the evolutionary history of birds-of-paradise has been studied with morphological data [1], molecular distances [4, 5], and a single mitochondrial gene [6], but the conclusions have been largely incongruent. The most comprehensive phylogenetic study at present includes all 41 species and is based on DNA-sequence data from both mitochondrial (cytochrome B) and nuclear genes (ornithine decarboxylase introns ODC6 and ODC7) [7]. This study suggests that the birds-of-paradise started to diverge during late Oligocene or early Miocene and could be divided into five main clades. The sexually monomorphic genera *Manucodia, Phonygammus*, and *Lycocorax* form a monophyletic clade (Clade A; Fig. 1 in Irestedt et al. 2009 [7]), which is suggested to be sister to the other four clades that include species with more or less strong sexual dimorphism (here referred to as “core birds-of-paradise”). Among the latter four clades, the genera *Pteridophora* and *Parotia* are suggested to form the earliest diverging clade (Clade B; Fig. 1 in Irestedt et al. 2009 [7]), followed by a clade consisting of the genera *Seleucidis, Drepanornis, Semioptera, Ptiloris*, and *Lophorina* (Clade C; Fig. 1 in Irestedt et al. 2009 [7]). The last two sister clades are formed by *Epimachus, Paradigalla*, and *Astrapia* (Clade D; Fig. 1 in Irestedt et al. 2009 [7]), and *Diphyllodes, Cicinnurus*, and *Paradisaea* (Clade E; Fig. 1 in Irestedt et al. 2009 [7]), respectively. In general, the phylogenetic hypothesis presented in Irestedt et al. (2009) [7] receives strong branch support (posterior probabilities), but several nodes are still weakly supported and there is incongruence among gene trees. Recently, Irestedt and colleagues [8] and Scholes and Laman [9] argued for the Superb birds-of-paradise to be split into several species, based on genetics, morphology and courtship behavior. Thus, while preliminary genetic analyses have outlined the major phylogenetic divisions, the interspecific relationships remain largely unresolved.

**Figure 1:**
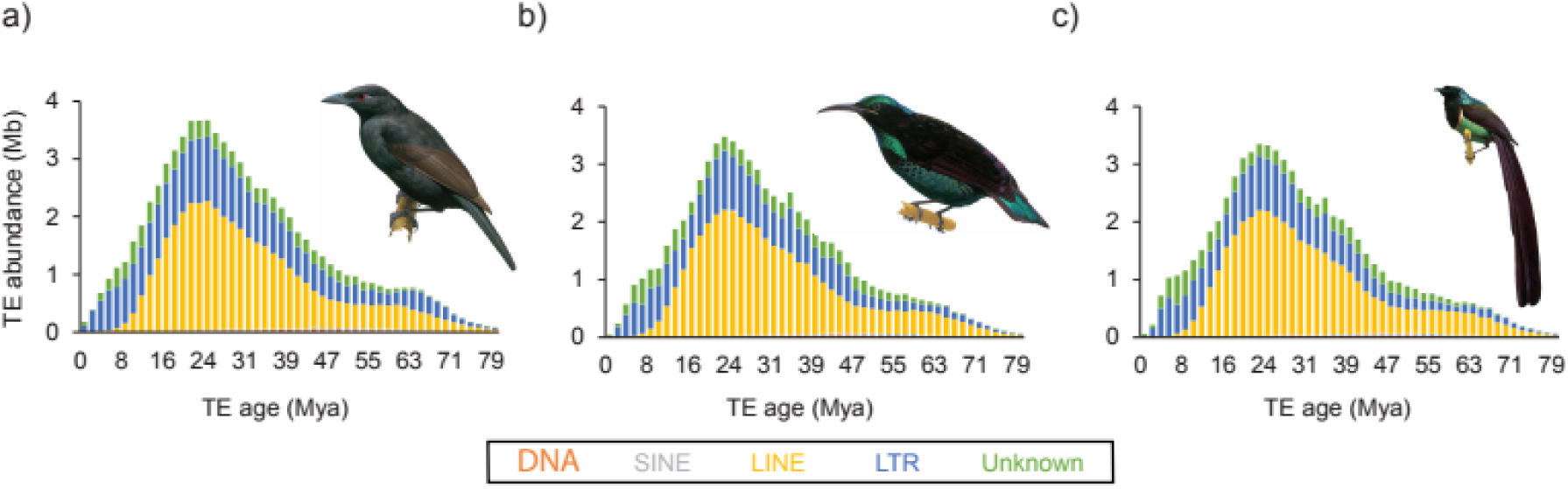
Repeat landscapes of *Astrapia rothschildi, Lycocorax pyrrhopterus*, and *Ptiloris paradisaeus*. Total amounts of TE-derived bp are plotted against relative age, approximated by per-copy Kimura 2-parameter distance to the TE consensus sequence and scaled using a fourfold degenerate mutation rate of Passeriformes of 3.175 substitutions/site/million years. TE families were grouped as “DNA transposons” (red), “SINEs” (grey), “LINEs” (yellow), “LTRs” (blue), and “Unknown” (green).

Birds-of-paradise are most widely known for their extravagant feather types, coloration and mating behaviors [3]. In addition, they also exhibit an array of bill shapes (often specialised on foraging behavior), and body morphologies and sizes [3]. Ornament feather types include ‘wire-type’ feathers (e.g. Twelve-wired bird-of-paradise (*Seleucidis melanoleuca*)), erectile head plumes (e.g. King of Saxony bird-of-paradise (*Pteridophora alberti*)), significantly elongated tail feathers (e.g. Ribbon-tailed Astrapia (*Astrapia mayeri*)) or finely-filamental flank plumes (e.g. Lesser bird-of-paradise (*Paradisaea minor*); see Frith and Bheeler 1998 [3]). Feathers and coloration are crucial components of their mating displays (see Discussion below). Polygynous birds-of-paradise show aggregated leks high in tree tops, less aggregated leks on lower levels or the forest floor (often exploded leks), and even solitary mating displays [3].

The array of extravagant phenotypes found in birds-of-paradise makes them an interesting model to study evolution. However, fresh tissue samples from birds-of-paradise are extremely limited and currently only about 50% of all species are represented in biobanks. Fortunately, the current revolution in sequencing technologies and laboratory methods does not only enable us to sequence whole-genome data from non-model organisms, but it also allows us to harvest genome information from specimens in museum collections [10]. Only recently, have these technological advances enabled researchers to investigate genome-wide signals of evolution using comparative and population genomic approaches in birds [11–14].

In the current study, we made use of these technological advances to generate *de novo* genomes for three birds-of-paradise species and re-sequenced the genomes of two other species from museum samples. Using these genomes, we were able to contrast the trajectory of genome evolution across passerines and simultaneously evaluate which genomic features have evolved during the radiation of birds-of-paradise. We identified a set of candidate genes that most likely have contributed to the extraordinary diversity in phenotypic traits found in birds-of-paradise.

## Results

### Assembly and gene annotation

We *de novo* assembled the genomes of *Lycocorax pyrrhopterus, Ptiloris paradiseus* and *Astrapia rothschildi* using paired-end and mate pair Illumina sequence data, and performed reference based mapping for *Pteridophora alberti* and *Paradisaea rubra*. Scaffold N50 ranged from 4.2 Mb (L. *pyrrhopterus*) to 7.7 Mb (*A. rothschildi*), and the number of scaffolds from 2,062 (*P. paradiseus*) to 3,216 (*L. pyrrhopterus;* Supplementary Table S1). All assemblies showed a genome size around 1 Gb. BUSCO2 [15] scores for complete genes (using the aves_odb9 database) found in the respective assemblies ranged from 93.8% to 95.1%, indicating a high completeness (Supplementary Table S2). Next we annotated the genomes using homology to proteins of closely related species as well as *de novo* gene prediction. Gene numbers ranged from 16,260 (*A. rothschildi*) to 17,269 (*P. paradiseus;* see Supplementary Table S3).

### Repeat evolution in birds-of-paradise

Our repeat annotation analyses (Supplementary Table S4) suggest that the genomes of birds-of-paradise contain repeat densities (~7%) and compositions (mostly chicken repeat 1 (CR1) long interspersed nuclear elements (LINEs), followed by retroviral long terminal repeats (LTRs)) well within the usual range of avian genomes [16]. However, we identified 16 novel LTR families (Supplementary Table S5) with no sequence similarity to each other or to LTR families known from in-depth annotations of chicken, zebra finch, and collared flycatcher [17, 18]. Interestingly, we find that activity of CR1 LINEs ceased recently in the three birds-of-paradise and was replaced by activity of retroviral LTRs (Fig. 1). The inferred timing of the TE (Tandem element) activity or accumulation peak (Fig. 1) corresponds to the radiation of birds-of-paradise (inferred in Irestedt et al. 2009 [7]). We also found that the genome assembly of *Lycocorax pyrrhopterus* exhibits slightly higher repeat densities than those of the two other birds-of-paradise (Supplementary Table S4) and slightly more recent TE activity (Fig. 1). A possible explanation for this is that this is the only female bird-of-paradise assembly, thus containing the female-specific W chromosome which is highly repetitive [16].

### Genome synteny to the collared flycatcher

We found strong synteny of the three *de novo* assembled birds-of-paradise genomes (*Lycocorax pyrrhopterus, Ptiloris paradiseus, Astrapia rothschildi*) to that of the collared flycatcher (Fig. 2 and Supplementary Figures S1–3). Only a few cases were found where scaffolds of the birds-of-paradise genomes mapped to different chromosomes in the collared flycatcher genome.

**Figure 2:**
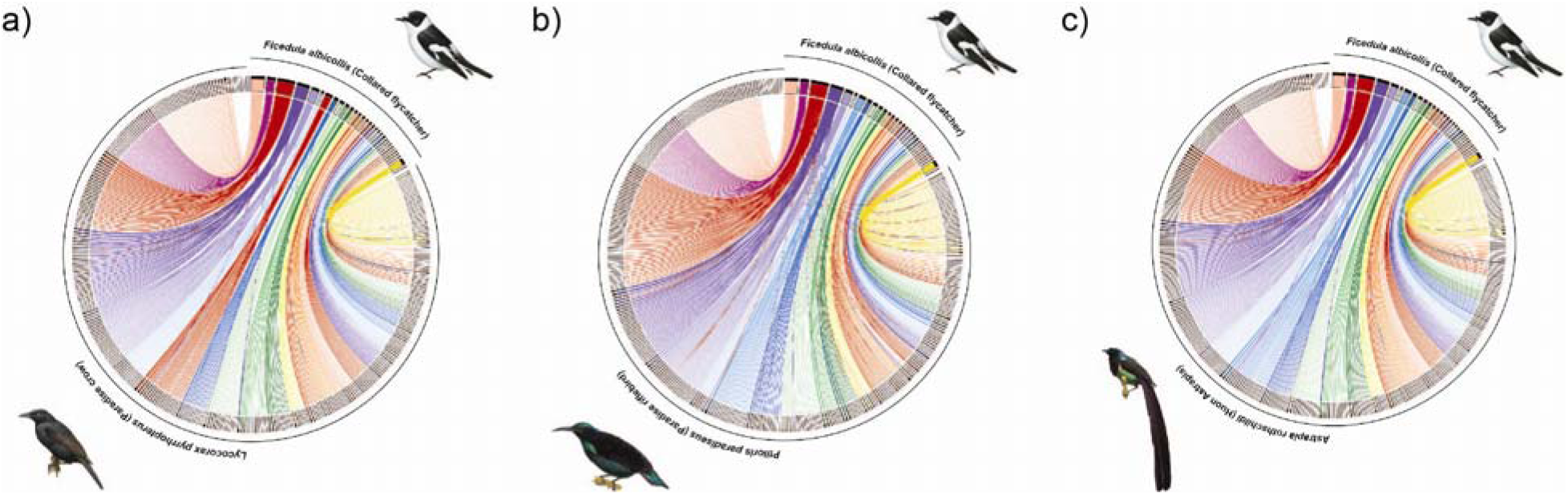
Chromosomal synteny plot between the collared flycatcher and (a) the paradise crow, (b) paradise riflebird and (c) the Huon Astrapia. The plot shows scaffolds bigger than 50kb and links (alignments) bigger than 2kb.

### Phylogeny

We found 4,656 one-to-one orthologous genes to be present in all eight sampled bird genomes (5 birds-of-paradise and 3 outgroup songbirds). A phylogeny inferred using these orthologs shows a topology with high bootstrap scores (Fig. 3 and Supplementary Figure S4). However, the sole use of bootstrapping or Bayesian posterior probabilities in analyses of large scale data sets has come into question in recent years [19]. Studies based on genome-wide data have shown that phylogenetic trees with full bootstrap or Bayesian posterior probability support can exhibit different topologies (e.g. Jarvis et al. 2014 [20] and Prum et al. 2015 [21]; discussed in Suh 2016 [19]). Thus, next we performed a concordance analysis by comparing gene trees for the 4,656 single-copy orthologs to the inferred species topology. We find strong concordance for the older splits in our phylogeny (see Supplementary Figure S4). However, the splits between Ptiloris and its sister clade, which contains Astrapia and Paradisaea, and the split between Astrapia and Paradisaea itself showed much lower concordance values, 0.31 and 0.26, respectively. Only ~10% of the gene trees exactly matched the topology of the inferred species tree and we find an average Robinson-Foulds distance of 3.92 for all gene trees compared to the species tree (Supplementary Table S6). A Robinson-Foulds distance of 0 would indicate that the two tree topologies (species to gene tree) are identical. The highest supported species tree topology (Fig. 3 and Supplementary Figure S4) is identical to the birds-of-paradise species tree constructed in Irestedt et al. 2009 [7]. Overall, we find that the birds-of-paradise form a monophyletic clade, with the crow (*Corvus cornix*) being the most closely related sister taxa, in most gene trees (74%). Within the birds-of-paradise clade, we further distinguish a core birds-of-paradise clade, which consist of 4 of the 5 species in our sample (excluding only the paradise crow, *Lycocorax pyrrhopterus*).

**Figure 3:**
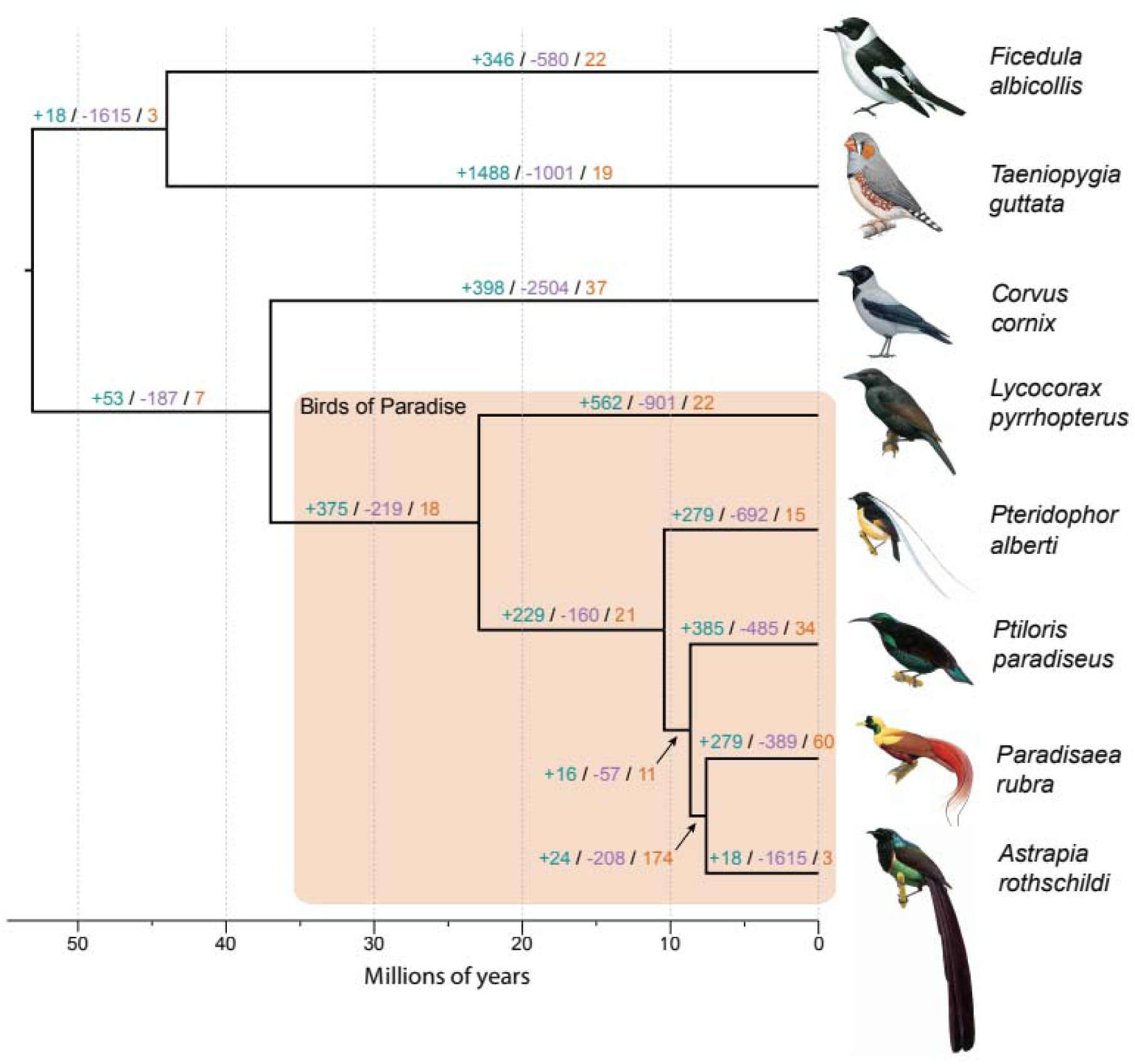
The birds-of-paradise phylogeny. The phylogenetic tree is based on ML and coalescent-based statistical binning of 4,656 genes, and scaled using the divergence times between crow and the birds-of-paradise, and zebra finch and flycatcher (obtained from Timetree.org) as calibration points. Branches are labeled as: # gene family expansions / # gene family contractions / # rapidly evolving gene families.

### Positive selection in the birds-of-paradise

We carried out positive selection analyses using all previously ascertained orthologous genes (8,134 genes present in at least seven out of the eight species) on the branch leading to the birds-of-paradise. First, we investigated saturation by calculating pairwise dN/dS ratios. The inferred values did not show any signs of saturation (Supplementary Table S7). To infer positive selection on the branch of the birds-of-paradise, we used the BUSTED model in HyPhy (similar to branch-sites model; [22]). We found 178 genes to be under selection (p value < 0.05; gene symbol annotation for 175 of the 178 genes can be found in Supplementary Table S8). GO enrichment resulted in 47 enriched GO terms using a 0.05 FDR cutoff (262 before correction; Supplementary Table S9). GO analysis showed enrichment of several categories related to collagen, skeletal and feather development, eye development, and glycogen synthesis and regulation (Supplementary Table S9).

### Gene gain and loss

We identified 9,012 gene families across all 8 species. Using CAFE [23] we inferred 98 rapidly evolving families within the birds-of-paradise clade. Supplementary Table S10 summarizes the gene family changes for all 8 species (also see Fig. 3). Zebra finch had the highest average expansion rate across all families at 0.0916, while the hooded crow had the lowest average expansion rate at -0.1724, meaning that they have the most gene family contractions. Gene gain loss rates can be found in Supplementary Table S11. Next, we tested for enrichment of GO terms in the set of families rapidly evolving in the birds-of-paradise clade. Gene families were assigned GO terms based off the Ensembl GO predictions for flycatcher and zebra finch. In all, we were able to annotate 6,350 gene families with at least one GO term. Using a Fisher’s exact test on the set of 98 rapidly evolving families in the birds-of-paradise, we find 25 enriched GO terms in 20 families (FDR 0.05; Supplementary Table S12). All the gene gain and loss results can be found online (https://cgi.soic.indiana.edu/~grthomas/cafe/bop/main.html).

## Discussion

Renowned for their extravagant plumage and elaborate courtship displays, the birds-of-paradise are among the most prominent examples of how sexual selection can give rise to extreme phenotypic diversity. Despite extensive work on systematics and a long-standing interest in the evolution of their different mating behaviors, the genomic changes that underlie this phenotypic radiation have received little attention. Here, we have assembled representative genomes for the five main birds-of-paradise clades and characterized differences in genome evolution within the family and relative to other avian groups. We reconstructed the main structure of the family phylogeny, uncovered substantial changes in the TE landscape and identified a list of genes under selection and gene families significantly expanded or contracted that are known to be involved with many phenotypic traits for which birds-of-paradise are renowned. Below, we discuss these different genomic features and how they might have contributed to the evolution of birds-of-paradise.

## Genome synteny and phylogeny

We found genome synteny (here in comparison to the collared flycatcher) to be highly conserved for all three *de novo* assembled genomes (Fig. 2 and Supplementary Figures S1–3). Only a few cases were recorded where regions of scaffolds of the birds-of-paradise genomes aligned to different chromosomes of the collared flycatcher. These could be artifacts of the genome assembly process or be caused by chromosomal fusions/fissions. Passerine birds show variable numbers of chromosomes (72–84 [16]). However, while passerines’ chromosome numbers do not vary as much as other groups’, such as Charadriiformes (shorebirds, 40–100 [16]), they still show frequent fissions and fusions of macro-and microchromosomes. Apart from these fission and fusion events, studies have shown a high degree of genome synteny even between Galloanseres (chicken) and Neognathae (approx. 80–90 mya divergence, reviewed in Ellegren 2010 [24]; Poelstra et al. 2014 [11]). However, genomes with higher continuity, generated with long-read technologies or using long-range scaffolding methods (such as HiC [25]), or a combination thereof, will be needed to get a more detailed view of rearrangements in genomes of birds-of-paradise.

Our analyses reconstructed a phylogenetic tree topology congruent with the one presented in Irestedt et al. 2009 [7] (Fig. 3 and Supplementary Figure 4). However, while bootstrapping found full support for the topology of the species tree, congruence analysis found high discordance for the two most recent branches (*Ptiloris* and its sister clade (*Astrapia* and *Paradisaea*) and the split between *Astrapia* and *Paradisaea*). Furthermore, we found the highest supported tree topology to be based on only 10% of all gene trees (Supplementary Table S6). This could be caused by incomplete lineage sorting (ILS), which refers to the persistence of ancestral polymorphisms across multiple speciation events [26]. Jarvis et al. 2014 [20] and Suh et al. 2015 [27] showed that ILS is a common phenomenon on short branches in the bird Tree of Life. Another possibility could be hybridization, a phenomenon frequently recorded in birds-of-paradise [3]. Overall, most gene tree topologies (74%) support the monophyly for the birds-of-paradise and the core birds-of-paradise clades.

### Repeats and their possible role in the evolution of birds-of-paradise

A growing body of literature is emerging that proposes an important role of TEs in speciation and evolution (see e.g. Feschotte 2008 [28], Oliver and Greene 2009 [29]). Bursts of TE activity are often lineage and species-specific, which highlights their potential role in speciation [30]. This is further supported by the fact that TE activity bursts often correlate with the speciation timing of the respective species or species group [31]. Similarly, we find a burst of TE activity within all three *de novo* assembled genomes (*Lycocorax pyrrhopterus, Ptiloris paradiseus, Astrapia rothschildi*) dating back to about 24 mya (Fig. 1). The timing fits the emergence and radiation of birds-of-paradise (see Fig 3). The fact that we found 16 novel families of retroviral LTRs suggests multiple recent germline invasions of the birds-of-paradise lineage by retroviruses. The recent cessation of activity of CR1 LINEs and instead recent activity of retroviral LTRs (Fig. 1) is in line with similar trends in collared flycatcher and hooded crow [16, 17]. This suggests that recent activity of retroviral LTRs might be a general genomic feature of songbirds, however, with different families of retroviruses being present and active in each songbird lineage. It is thus likely that the diversification of birds-of-paradise was influenced by lineage specificity of their TE repertoires through retroviral germline invasions and smaller activity bursts.

### Coloration, feather and skeletal development in birds-of-paradise

The diverse array of color patterns exhibited by birds-of-paradise involve both pigmentary and structural coloration mechanisms. Coloration via pigmentation is achieved by pigment absorption of diffusely scattered light in a specific wavelength range. Pigments such as carotenoids are frequently associated with red and yellow hues in birds, whereas light absorption by various classes of melanin give rise to black plumage features common to many birds-of paradise [32]. On the other hand, structural coloration is caused by light reflection of quasiordered spongy structures of the feather barbs and melanosomes in feather barbules [33, 34]. The plumages of male birds-of-paradise feature both coloration types to various degrees, and some species such as the Lawes’ parotia (*Parotia lawesii*) use angular-dependent spectral color shifts of their structural feathers in their elaborate display rituals to attract females [35, 36]. Most core birds-of-paradise show a strong sexual dimorphism, with highly ornamented males and reduced ornamentation in females [3]. Dale and colleagues recently showed that sexual selection on male ornamentation in birds has antagonistic effects, where male coloration is increasing, while females show a strong reduction in coloration [37]. This is very apparent in polygynous core birds-of-paradise, where females between species and sometimes even between genera look highly similar. Given their strong sexual dimorphism and its important role in mating success, we would expect genes important for coloration, morphology and feather structure to be under strong selection in the birds-of-paradise. In accordance with this prediction, we found several GO categories enriched in positively selected genes in birds-of-paradise that could be associated with these phenotypes.

One such gene is ADAMTS20, which is crucial for melanocyte development. ADAMTS20 has been shown to cause white belt formation in the lumbar region of mice [38]. Nonsense or missense mutations in this gene disrupt the function of KIT, a protein that regulates pigment cell development [38]. In mammals and birds pigment patterns are exclusively produced by melanocytes. Thus, this gene could be a strong candidate for differential coloration in the birds-of-paradise. Another gene under positive selection with a potential role in coloration is ATP7B. It is a copper-transporting P-type ATPase and thought to translate into a melanosomal protein (see Bennett and Lamoreux 2003 [39] for a review). Copper is crucial for melanin synthesis because tyrosinase contains copper and thus ATP7B might play a crucial role in pigment formation.

Genes in GO categories involving collagen and the extracellular matrix are likely affecting morphology (feather, craniofacial and skeletal muscle development) in birds-of-paradise (Supplementary Table S9). Several genes under positive selection fall into these GO categories. FGFR1 (Fibroblast growth factor receptor 1) is implicated in feather development [40]. In humans it has further been shown to be involved in several diseases associated with craniofacial defects (OMIM; http://www.ncbi.nlm.nih.gov/omim/). ALDH3A2 (aldehyde dehydrogenase 3 family member A2), a membrane-associated protein and SPECC1L are implicated in craniofacial disorders (e.g. Van Der Woude Syndrome) in humans [41] and ALDH3A2 has been listed as a candidate gene for beak development in birds [12]. GAB2 (GRB2-associated-binding protein 2) is an important gene in osteoclastogenesis and bone homeostasis [42] (bone remodeling). PAPSS2 (Bifunctional 3'-phosphoadenosine 5'-phosphosulfate synthetase 2) plays an important role in cartilage development [43]. Similarly, DCST2 (*DC-STAMP domain containing 2*) is an important regulator of osteoclast cell-fusion in bone homeostasis [44], and has been shown to be associated with body length in early life and height in adulthood [45]. MYF5 (Myogenic factor 5) has been shown to be important for skeletal muscle phenotype and initiates the myogenic program [46] (muscle tissue formation). APOBEC2 seems to play a role in muscle development (skeletal and heart muscle) in chickens [47].

### APOBECs and their potential role in the immune system

Intriguingly, APOBECs have also been shown to have important functions in the immune systems of vertebrates, where they act as restriction factors in the defense against a range of retroviruses and retrotransposons [48, 49]. Functioning as cytosine deaminases they act against endogenous retroviruses (ERVs), especially Long terminal repeat retrotransposons (LTR) by interfering with the reverse transcription and by hypermutating retrotransposon DNA. A recent study on 123 vertebrates showed that birds have the strongest hypermutation signals, especially oscine passerines [50] (such as zebra finch and medium ground finch). This study also demonstrated that edited retrotransposons may preferentially be retained in active regions of the genome, such as exons, promoters, etc. (hypermutation decreases their potential for mobility). Thus, it seems very likely that retrotransposon editing via APOBECs has an important role in the innate immunity of vertebrates as well as in genome evolution. Congruently, we found a burst in recent activity of retroviral LTRs in the genomes of birds-of-paradise, a similar signal was further found in other passerines [16, 17]. This could also explain the why we found APOBEC2 to be under positive selection.

In concert, the inferred genes under positive selection and the results of the GO category enrichment analyses indicate that positive selection has played a role in shaping morphological phenotypes of the birds-of-paradise and targeted developmental genetics studies may further elucidate their specific roles in this family.

### Sensory system in the birds-of-paradise

#### Visual system

We also found two GO categories associated with eye development and function to be enriched for positively selected genes in birds-of-paradise, namely "retina development in camera-type eye” and “photoreceptor outer segment”. Genes that showed positive selection and are known to have critical roles in eye function and development include CABP4, NR2E3, IMPG1, GNB1, AKAP13, MGARP, CDADC1 and MYOC. For example, CaBP4 is a member of a subfamily of calmodulin-like neuronal Ca2Þ-binding proteins (CaBP1–8) and is essential for normal photoreceptor synaptic function via continuous release of neurotransmitter in retinal photoreceptor cells [51]. NR2E3, a photoreceptor-specific nuclear receptor is a transcription factor important for retinal development [52]. Transcription analysis of MYOC indicates a structural or functional role of myocilin in the regulation of aqueous humor outflow that may influence intraocular pressure, and in the optical nerve [53].

There are no single obvious explanations for selection on vision in birds-of-paradise. Evidence for co-evolution between coloration and vision in birds is weak (see e.g. Lind et al. 2017 [54], Price 2017 [55], but see Mundy et al. 2016 [56] and Bloch 2015 [57]). Another phenotype that might be associated with selection on vision is the diverse array of mating displays in some core birds-of-paradise. Many species, such as the Lawes’ parotia (*Parotia lawesii*) modify color by changing the angle of the light reflection [35, 36], which requires the visual system to be able to process the fine nuances of these color changes. However, the fact that (color) vision serves many purposes (including e.g. foraging, etc.) makes it very difficult to establish co-evolution between coloration and color vision [54]. We can thus only speculate at this point about the potential role of coloration or mating displays in the selection of vision genes found in birds-of-paradise.

#### Olfactory system

Another often overlooked sensory system in birds is odor perception. Olfactory receptors (ORs) are important in odor perception and detection of chemical cues. In many animal taxa, including birds, it has been shown that olfaction is crucial to identify species [58], relatedness [59], individuals [60], as well as for mate choice [61] and in foraging [62]. In concordance with previous studies we found this gene family to expand rapidly in the zebra finch [63]. Even more so, the zebra finch showed the strongest expansion (+17 genes). Furthermore, we find a rapid expansion on the branch leading to the core birds-of-paradise (+5 genes) and further in *Astrapia* (+6 genes). Interestingly, olfactory receptor genes show rapid contractions in the paradise crow (-6 genes), the hooded crow (-9 genes) and the collared flycatcher (-5 genes). This is in line with a study that suggested poor olfactory development in the Japanese jungle crow (*Corvus macrorhynchos*) [64]. Olfactory could serve many functions in birds-of-paradise e.g. in species recognition (to avoid extensive hybridization), individual recognition, mating or foraging (given their extensive diet breadth), among others.

#### Startle response and adult locomotory behavior

Startle response is an important behavioral trait. It is the ability to quickly react to the presence of a stimuli, such as the presence of predators. It could be crucial for core birds-of-paradise that show extravagant lekking behavior, arguably especially for those taxa that congregate at large leks at highly visible places. Being highly visible means that they need to be able to look out for predators and react to them quickly, and indeed Frith & Beehler 1998 [3] mention that lekking birds-of-paradise appear to be constantly on the lookout for predators. We find a gene family associated with startle response and adult locomotory behavior to be evolving significantly faster than under a neutral model on the branch leading the core birds-of-paradise (+5 genes). It is even further expanded in the genus *Paradisaea* (+3 genes). Interestingly, species of the genus *Paradisaea* have aggregated leks high in emergent trees and thus may be more visible to the numerous birds of prey that inhabit the region, while most other core birds-of-paradise display on lower levels in trees or on the forest floor and have less aggregated leks (exploded lek) or solitary display [3]. This gene family is contracted in the two outgroups, the zebra finch (-4 genes) and the collared flycatcher (-2 genes), as well as the monochromatic, non-lekking Paradise crow (-1 genes). We found no expansion or contraction in the hooded crow genome.

#### Other positively selected genes and enriched GO categories

Other interesting genes under positive selection include CTSD (Cathepsin D), a gene that has been shown to play a key (enzymatic) role in yolk production in chicken [65]. CTSD is primarily important for egg yolk and egg weight [65, 66]. We also found several genes important for sexual development to be under positive selection. These include CBX2, SPAG16, TAF4B, SPATA5L1 and DCST2. CBX2 (Chromobox homolog 2) has been shown to determine sex in humans, maybe even more so than X/Y chromosomes [67]. It is essential for the expression of SRY (sex determining region on the Y chromosome), which determines sex in most eutherian mammals [68]. Other genes include SPAG16 (Sperm-associated antigen 16) and STRA8 (Stimulated By Retinoic Acid 8), both of which are essential for spermatogenesis [69], and TAF4B (Transcription initiation factor TFIID subunit 4B), which is important for healthy ovarian aging and female fertility in mice and humans [70].

Interestingly, we found several GO categories related to glycogen synthesis and regulation to be enriched in the set of positively selected genes in birds-of-paradise (Supplementary Table S9). Genes under positive selection important for glycogen synthesis and regulation include SLC5A9, G6PC2, AGL, B3GLCT, PHLDA3 and IDE. SLC5A9 (Solute Carrier Family 5 Member 9), also called SGLT4, is a glucose transporter [71]. G6PC2 (Glucose-6-phosphatase 2) is involved in catalyzing the hydrolysis of glucose-6-phosphate, the terminal step in gluconeogenic and glycogenolytic pathways, which allow glucose to be released into the bloodstream [72]. AGL (amylo-1,6-glucosidase) is a glycogen debranching enzyme^99^, which facilitates the_break-down of glycogen (storage of glycogen in the body). We also found IDE (insulin degrading enzyme) to be under positive selection in birds-of-paradise. This gene is a large zinc-binding protease and degrades the B chain of insulin [73]. In birds, glucose is utilized in a variety of functions, with the main one being energy production through cellular oxidation, glycogen synthesis, etc. (see Braun and [74] 2008 for a review). Interestingly, birds maintain higher levels of plasma glucose than other vertebrates of similar body mass, but seem to store very little as glycogen [74]. On the contrary to other vertebrates, plasma glucose levels are insensitive to insulin in birds (see e.g. Sweazea et al. 2006 [74]). It thus seems surprising that we found a significant enrichment of positively selected genes involved in 'positive regulation of glucose import in response to insulin stimulus' in birds-of-paradise. On the other hand, it appears that gluconeogenesis plays an important role in maintenance of plasma glucose levels in birds [75]. Furthermore, it has been shown that in some birds, such as pigeons, blood glucose levels are significantly increased during courtship and mating, a time of significant energy requirement [76].

High plasma glucose levels, high metabolic rates and high body temperatures should increase oxidative stress in birds [77]. However, birds have developed efficient mechanisms to prevent tissue damage of oxidative stress (see e.g. Holmes et al. 2001 [77]). Furthermore, studies have shown that mitochondria in different bird tissues produce much lower levels of reactive oxygen [78]. In addition, they show higher levels of antioxidants superoxide dismutase, catalase and glutathione peroxidase [78]. Intriguingly, we found a significant GO enrichment of 'glutathione peroxidase activity' in the set of positively selected genes in birds-of-paradise.

Beside energy storage, glycogen seems to have a function in the visual system of some birds. Several studies have shown a high concentration of glycogen B in lenses of flying birds [79]. The function of glycogen in lenses of flying birds is unknown, but a structural function, more specifically the maintenance of the refractive index of the lens, is suspected [79]. Castillo and colleagues were able to detect high concentrations of glycogen in pigeon and dove, but not the chicken [79] (which they classify as a ground-running bird).

## Conclusions

We found several genes with known function in coloration, feather, and skeletal development to be under positive selection in birds-of-paradise. This is in accordance with our prediction that phenotypic evolution in birds-of-paradise should have left strong genomic signatures. Furthermore, positively selected genes were enriched for GO terms associated with collagen and extracellular matrix development. While these gene categories all are obvious candidates for being important in the evolution of birds-of-paradise’s phenotypic and behavioral diversity, we also found enrichment in other positively selected genes that are not as straight forward to explain. These include eye development and function, glycogen synthesis and regulation, and glutathione peroxidase activity. Gene gain loss analyses further revealed significant expansions in gene families associated with ‘startle response’ (response to predators) and olfactory function. On the genome level, we found indications of a highly conserved synteny between birds-of-paradise and other passerine birds, such as the collared flycatcher. Similar to other passerine groups we also found strong signatures of recent activity of novel retroviral LTRs in the genomes. Birds-of-paradise show positive selection on APOBEC2, likely to counteract deleterious effects of LTRs, by decreasing their activity.

Although recent advances in documenting the phenotypic and behavioral diversity in the birds-of-paradise continues to generate intense interest in this model system, we still have very limited understanding of the processes that have shaped their evolution. Here, we provide a first glimpse into genomic features underlying the diverse array of phenotypes found in birds-of-paradise. However, more in depth analyses will be needed to verify a causal relationship between signatures of selection in the birds-of-paradise genome and the unique diversity of phenotypic traits, or to investigate genome structure changes with higher resolution. Fortunately, technologies keep advancing, and along with decreasing costs for sequencing, we will soon be able to gain more information about this fascinating, but understudied family of birds.

## Data description and Analyses

### Sampling and DNA extraction

For the three *de novo* genome assemblies, *Lycocorax pyrrhopterus* (ZMUC149607; collected 2013, Obi Island, Indonesia), *Ptiloris paradiseus* (ANWC43271; collected 1990, New South Wales, Australia), and *Astrapia rothschildi* (KU Birds 93602; collected 2001, Morobe Province, Papua New Guinea) DNA was extracted from fresh tissue samples using the Qiagen QIAamp DNA Mini Kit according to the manufacturer's instructions. The *de novo* libraries with different insert sizes (see below) were prepared by Science for Life Laboratory, Stockholm. For the two re-sequenced genomes, *Pteridophora alberti* (NRM571458; collected 1951, Eastern Range, New Guinea) and *Paradisaea rubra* (NRM700233; collected 1949, Batanta Island, New Guinea), we sampled footpads and extracted DNA using the Qiagen QIAamp DNA Micro Kit to the manufacturer's instructions. We applied precautions for working with museum samples described in Irestedt et al., (2006) [80]. Sequencing libraries for these two samples were prepared using the protocol published by Meyer and Kircher, (2010) [81]. This method was specifically developed to generate sequencing libraries for low input DNA, showing DNA damage typical for museum or ancient samples.

### Genome Sequencing, Assembly, and Quality Assessment

We prepared two paired-end (overlapping and 450bp average insert size) and two mate pair libraries (3kb and 8kb average insert size) for each of the three *de novo* assemblies (*Ptiloris paradiseus, Astrapia rothschildi*, and *Lycocorax pyrrhopterus*). All libraries, for the *de novo* and the reference-based mapping approaches were sequenced on a HiSeq2500 v4 at SciLife Stockholm, Sweden. We generated 2 lanes of sequencing for each *de novo* assembly and pooled the two reference-based samples on one lane. We first assessed the read qualities for all species using the program FastQC ^[82]^. For the three species, *Ptiloris paradiseus, Astrapia rothschildi*, and *Lycocorax pyrrhopterus* we then used the *preqc* ^[83]^ function of the SGA ^[84]^ assembler to (1) estimate the predicted genome size, (2) find the predicted correlation between k-mer sizes and N50 contig length and (3) assess different error statistics implemented in *preqc*. For *Ptiloris paradiseus, Astrapia rothschildi*, and *Lycocorax pyrrhopterus* reads were assembled using Allpaths-LG ^[85]^. To improve the assemblies, especially in repeat regions, GapCloser (part of the SOAPdenovo package ^[86]^) was used to fill in gaps in the assembly. Assemblies were then compared using CEGMA ^[87]^ and BUSCO2 ^[15]^. We added BUSCO2 scores for better comparisons at a later stage of the project. For the reference-based mapping, we mapped all reads back to the *Ptiloris paradiseus* assembly using BWA ^[88]^ (mem option), the resulting sam file was then processed using samtools ^[89]^. To do so, we first converted the sam file generated by BWA to the bam format, then sorted and indexed the file. Next we removed duplicates using Picard (http://broadinstitute.github.io/picard/) and realigned reads around indels using GATK ^[90]^. The consensus sequence for each of the two genomes was then called using ANGSD ^[91]^ (using the option -doFasta 3). [Genomes will be deposited in GenBank]

### Repeat Annotation

We predicted lineage-specific repetitive elements *de novo* in each of the three birds-of-paradise genome assemblies using RepeatModeler v. 1.0.8 ^[92]^. RepeatModeler constructs consensus sequences of repeats via the three complementary programs RECON ^[93]^, RepeatScout ^[94]^, and Tandem Repeats Finder ^[95]^. Next, we merged the resultant libraries with existing avian repeat consensus sequences from Repbase ^[96]^ (mostly from chicken and zebra finch), and recent indepth repeat annotations of collared flycatcher [97, 98] and hooded crow ^[99]^. Redundancies among the three birds-of-paradise libraries, and between these and existing avian repeats were removed using the ReannTE_mergeFasta.pl script (https://github.com/4ureliek/ReannTE/). For *Lycocorax pyrrhopterus* repeats, we manually inspected the RepeatModeler library of consensus sequences for reasons reviewed in Platt et al. (2016) ^[100]^ and because *Lycocorax pyrrhopterus* was the most repeat-rich genome among the three birds-of-paradise. Manual curation was performed using standard procedures [27, 101]’ namely screening of each repeat candidate against the *Lycocorax pyrrhopterus* assembly using blastn ^[102]^, extracting the 20 best hits including 2-kb flanks, and alignment of these per-candidate BLAST hits to the respective consensus sequence using MAFFT v. 6 [103]. Each alignment was inspected by eye and curated majority-rule consensus sequences were generated manually considering repeat boundaries and target site duplication motifs. This led to the identification of 33 long terminal repeat (LTR) retrotransposon consensus sequences (including 16 novel LTR families named as ‘lycPyrLTR*’) and three unclassified repeat consensus sequences (Supplementary Table S5). We then used this manually curated repeat library of *Lycocorax pyrrhopterus* to update the aforementioned merged library of avian and birds-of-paradise repeat consensus sequences. Subsequently, all three birds-of-paradise genome assemblies were annotated via RepeatMasker v. 4.0.6 and ‘-e ncbi’ ^[104]^ using this specific library (Supplementary Table S4). Landscapes of relative TE activity (i.e., the amount of TE-derived bp plotted against Kimura 2-parameter distance to respective TE consensus) were generated using the calcDivergenceFromAlign.pl and createRepeatLandscape.pl scripts of the RepeatMasker packages. To enhance plot readability, TE families were grouped into the subclasses "DNA transposon”, “SINE”, “LINE”, “LTR”, and “Unknown” (Fig. 1). We scaled the kimura substitution level with the four-fold degenerate mutation rate for Passeriformes (mean of 3.175 substitutions/site/million years for passerines sampled in Zhang et al. 2014 ^[13]^) to obtain an estimate of the timing of the inferred repeat activity in million years (mya).

### Genome Synteny

We also inferred genome architecture changes (synteny) between our three *de novo* assembled genomes and the chromosome-level assembly of the collared flycatcher. To do so, we first performed pairwise alignments using Satsuma ^[105]^ and then plotted the synteny using Circos plots ^[106]^. More precisely, we first performed asynchronous ‘battleship’-like local alignments using *SatsumaSynteny* to allow for time efficient pairwise alignments of the complete genomes. In order to avoid signals from tandem elements we used masked assemblies for the alignments. Synteny between genomes was then plotted using Circos and in-house perl scripts.

### Gene Annotation

We masked repeats (only tandem elements) in the genome prior to gene annotation. Contrary to the repeat annotation step, we did not mask simple repeats in this approach. Those were later soft-masked as part of the gene annotation pipeline Maker2 ^[107]^, to allow for more efficient mapping during gene annotation.

Gene annotation was performed using *ab-initio* gene prediction and homology-based gene annotation. To do so we used the genome annotation pipeline Maker2 ^[107]^, which is able to perform all the aforementioned genome annotation strategies. Previously published protein evidence (genome annotations) from Zhang et al. 2014 ^[13]^ were used for the homology-based gene prediction. To improve the genome annotation we used CEGMA to train the *ab-initio* gene predictor SNAP ^[108]^ before running Maker2 ^[107]^. We did not train the *de novo* gene predictor Augustus ^[109]^ because no training data set for birds was available.

### Ortholog Gene Calling

In the next step we inferred orthologous genes using PoFF ^[110]^. We included all five birds-of-paradise, as well as the hooded crow, the zebra finch and the collared flycatcher as outgroups. We ran PoFF using both the transcript files (in fasta format) and the transcript coordinates file (in gff3 format). The gff files were used (flag–*synteny*) to calculate the distances between paralogous genes to accurately distinguish between orthologous and paralogous genes. We then extracted the sequences for all one-to-one orthologs using a custom python script.

Next, we determined the number of genes with missing data in order to maximize the number of genes included in the subsequent analyses. For a gene to be included in our analyses, it had to be present in at least 75% of all species (7 out of 8 species), which resulted in a set of 8,134 genes. In order to minimize false positives in the subsequent positive selection analysis caused by alignment errors, we used the codon-based alignment algorithm of Prank ^[111]^ and further masked sites with possible alignment issues using Aliscore ^[112]^. Aliscore uses Monte Carlo resampling within sliding windows to identify low-quality alignments in amino acid alignments (converted by the program). The identified potential alignment issues were then removed from the nucleotide alignments using ALICUT ^[113]^.

### Intron Calling

In addition to exons, we also extracted intron information for the birds-of-paradise genomes (see Supplementary Table S3). To do so we used the extract_intron_gff3_from_gff3.py script (https://github.com/irusri/Extract-intron-from-gff3) to include intron coordinates into the gff file. We then parsed out all intron coordinates and extracted the intron sequences from the genomes using the exttract_seq_from_gff3.pl script (https://github.com/irusri/Extract-intron-from-gff3). All introns for the same gene were then concatenated using a custom python script.

### Phylogenetic analysis

The individual alignment files (we used exon sets without missing species, which resulted in 4,656 alignments) were then (1) converted to the phylip format individually, and (2) 200 randomly selected exon alignments were concatenated and then converted to phylip format using the catfasta2phyml.pl script (https://github.com/nylander/catfasta2phyml). We used the individual exon phylip files for gene tree reconstruction using RaxML ^[114]^ (using a GTR + G model). Subsequently, we binned the gene trees into a species tree and carried out bootstrapping using Astral ^[115, 116]^. Astral applies a statistical binning approach to combine similar gene trees, based on an incompatibility graph between gene trees and then chooses the most likely species tree under the multi-species coalescent model. Astral does not provide branch lengths needed for calibrating phylogenetic trees (used for the gene gain/loss analysis). So, we subsampled our data, and constructed a ML tree based on 200 randomly chosen and concatenated exons using ExaML ^[117]^. We then calibrated the species tree using the obtained branch lengths along with calibration points obtained from timetree.org using r8s ^[118]^. These calibration points are the estimated 44mya divergence time between flycatcher and zebra finch and the 37mya divergence time between crow and the birds-of-paradise.

Next we performed a concordance analysis. First, we rooted the gene trees based on the outgroup (Flycatcher, zebra finch). Then, for each node in the species tree we counted the number of gene trees that contained that node and divided that by the total number of gene trees. We next counted the number of gene trees that support a given topology (see Supplementary Table S6) and further calculated the Robison-Foulds distance between gene trees using RaxML.

### Inference of Positive selection

We inferred genes under positive selection using *dN/dS* ratios of 8,133 orthologs. First, we investigated saturation of synonymous sites in the phylogenetic sampling using pairwise comparisons in CodeML ^[119]^. The pairwise runmode of CodeML estimates *dN* and *dS* ratios using a ML approach between each species pair (Supplementary Table S7). We then investigated positive selection on the branch to the birds-of-paradise using the BUSTED model [22] (branch-site model) implemented in HyPhy ^[120]^. The branch-site test allows for inference of positive selection in specific branches (foreground branches) compared to the rest of the phylogeny (background branches). The significance of the model comparisons was determined using likelihood-ratio tests (LRT). We did not perform multiple testing corrections, such as Bonferroni correction, due to the fact that branch-site tests result in an access of non-significant p-values, which violates the assumption of a uniform distribution in multiple testing correction methods such as the Bonferroni correction. The genes were then assigned gene symbols. To do so, we first extracted all the respective zebra finch or collared flycatcher GeneBank protein accessions. We then converted the accessions into gene symbols using the online conversion tool bioDBnet ^[121]^. GO terms were obtained for the flycatcher assembly and assigned to orthologs that had a corresponding flycatcher transcript ID in Ensembl (7,305 genes out of 8,133). To determine enriched GO categories in positively selected genes, GO terms in genes inferred to have undergone positive selection were then compared to GO terms in all genes (with a GO term) using Fisher’s exact test with a false discovery rate cut-off of 0.05. We found 262 GO terms enriched in positively selected genes before FDR correction and 47 enriched after.

### Gene Gain-Loss

In order to identify rapidly evolving gene families in the birds-of-paradise we used the peptide annotations from all five birds-of-paradise species, along with the three outgroup species in our analysis: *Corvus cornix* (crow), *Taeniopygia guttata* (zebra finch), *Ficedula albicus* (flycatcher). The crow genes were obtained from NCBI and the zebra finch and flycatcher genes were acquired from ENSEMBL 86 ^[122]^. To ensure that each gene was counted only once, we used only the longest isoform of each protein in each species. We then performed an all-vs-all BLAST ^[102]^ search on these filtered sequences. The resulting e-values from the search were used as the main clustering criterion for the MCL program to group peptides into gene families ^[123]^. This resulted in 13,289 clusters. We then removed all clusters only present in a single species, resulting in 9,012 gene families. Since CAFE requires an ultrametric time tree as input, we used r8s to smooth the phylogenetic tree with calibration points based on the divergence time of crow and the birds-of-paradise at 37mya and of flycatcher and zebra finch at 44mya ^[124]^.

With the gene family data and ultrametric phylogeny (Figure 3) as input, we estimated gene gain and loss rates (λ) with CAFE v3.0 ^[125]^. This version of CAFE is able to estimate the amount of assembly and annotation error (*ε*) present in the input data using a distribution across the observed gene family counts and a pseudo-likelihood search. CAFE is then able to correct for this error and obtain a more accurate estimate of λ. We find an *ε* of about 0.01, which implies that 3% of gene families have observed counts that are not equal to their true counts. After correcting for this error rate, we find λ = 0.0021. This value for λ is considerably higher than those reported for other distantly related groups (Supplementary Table S11). GO terms were assigned to genes within families based on flycatcher and zebra finch gene IDs from Ensembl. We used these GO assignments to determine molecular functions that may be enriched in gene families that are rapidly evolving along the ancestral BOP lineage (Node BOP11 in Figure S1). GO terms in genes in families that are rapidly evolving along the BOP lineage were compared to all other GO terms using a Fisher’s exact test (FDR cut-off of 0.05). We found 36 genes in 26 families to have enriched GO terms before FDR correction and 25 genes in 20 families after.

## DATA AND SOFTWARE AVAILABILITY

All genomes will be deposited in GigaDB, and raw read data are currently deposited on NCBI (SRA archive). All results of the gene gain-loss analyses can be found online (https://cgi.soic.indiana.edu/~grthomas/cafe/bop/main.html).

## Acknowledgement

We thank Australian National Wildlife Collection, CSIRO Sustainable Ecosystems (Leo Joseph) for tissue sample from *Ptiloris paradiseus*, Natural History Museum of Denmark, University of Copenhagen (Knud Jønsson and Jan Bolding) for tissue samples from *Lycocorax pyrrhopterus*, and Natural History Museum, and Biodiversity Institute, University of Kansas (Robert Moyle) for tissue samples of *Astrapia rothschildi*. Daniel Osorio for helpful discussion of coloration and color vision in birds, and Nagarjun Vijay for help with the circos plots. MI and PGPE were supported by the Swedish Research Council (grant number 621–2013–5161 to PGPE and grant number 621–2014–5113 to MI). We also acknowledge support from Science for Life Laboratory, the National Genomics Infrastructure (NGI), Uppmax and the EvoLab at the University of California Berkeley for providing resources for massive parallel sequencing and computational infrastructure. The funders had no role in study design, data collection and analysis, decision to publish, or preparation of the manuscript.

## Supplementary Supplementary Figures

**Supplementary Figure S1:**
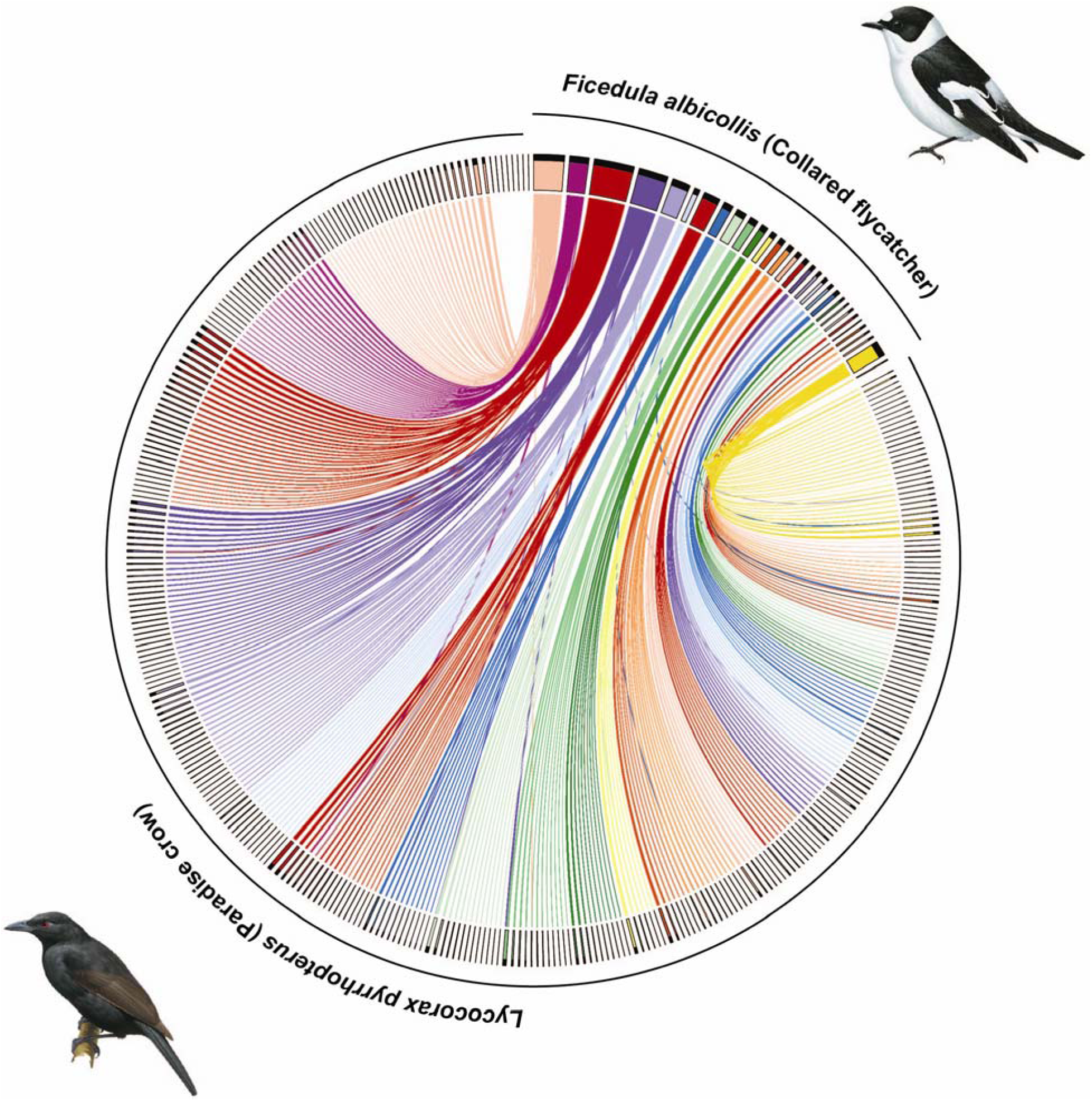
Chromosomal synteny plot between the collared flycatcher and the Paradise crow. The plot shows scaffolds bigger than 50kb and links (alignments) bigger than 2kb.

**Supplementary Figure S2:**
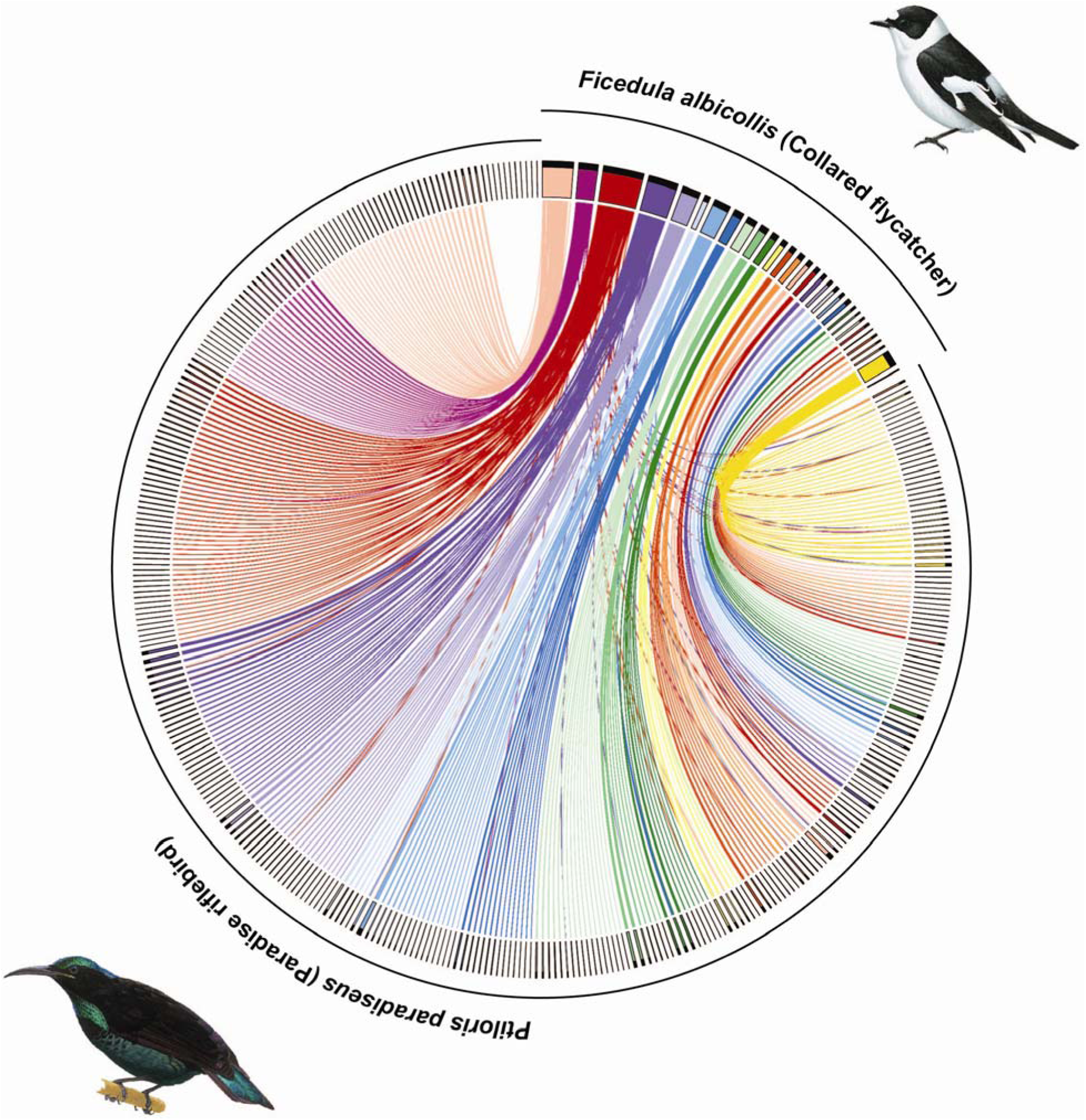
Chromosomal synteny plot between the collared flycatcher and the Paradise riflebird. The plot shows scaffolds bigger than 50kb and links (alignments) bigger than 2kb.

**Supplementary Figure S3:**
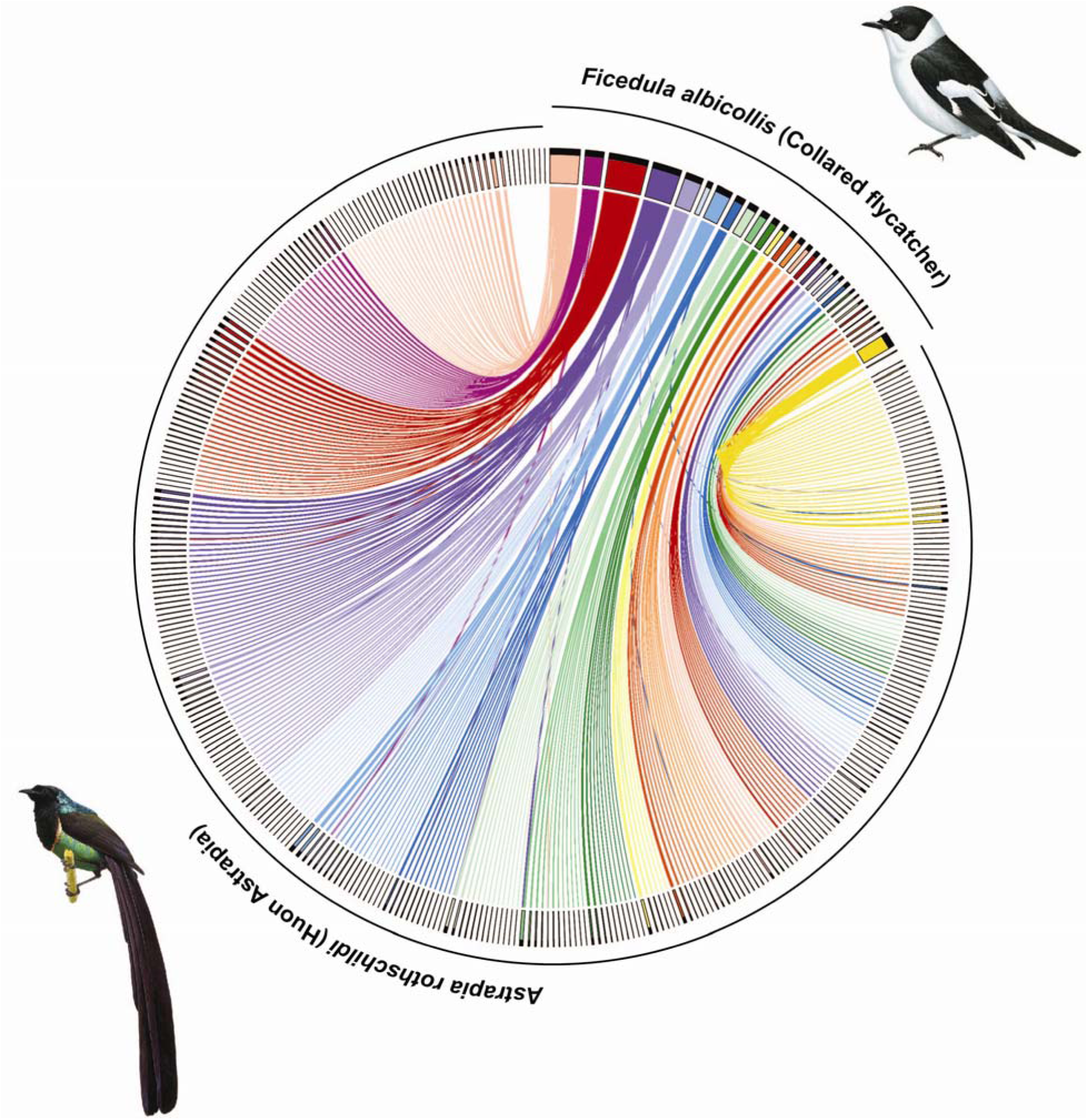
Chromosomal synteny plot between the collared flycatcher and the Huon Astrapia. The plot shows scaffolds bigger than 50kb and links (alignments) bigger than 2kb.

**Supplementary Figure S4:**
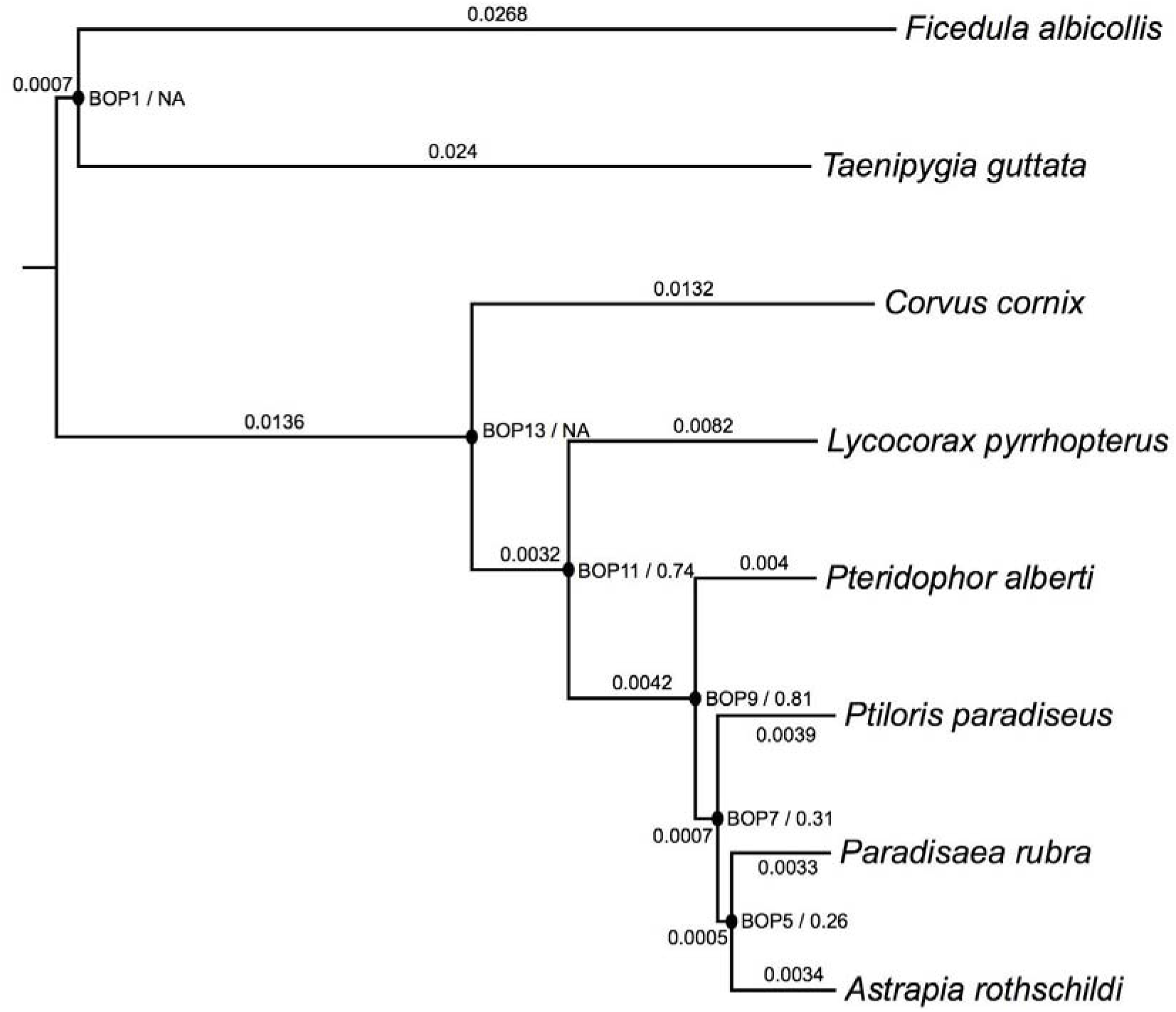
Phylogenetic species tree. The species tree was reconstructed from individual maximum likelihood-based gene trees using 4,656 exons and coalescent-based statistical binning (Astral). Branch lengths are depicted on the branches (calculated via a ML tree constructed using ExaML and 200 randomly selected genes). Nodes are labelled and concordance factor is shown next to the node labels (ie [node label] / [concordance factor]). All nodes have 100 bootstrap support.

## Supplementary Tables

**Supplementary Table S1:**
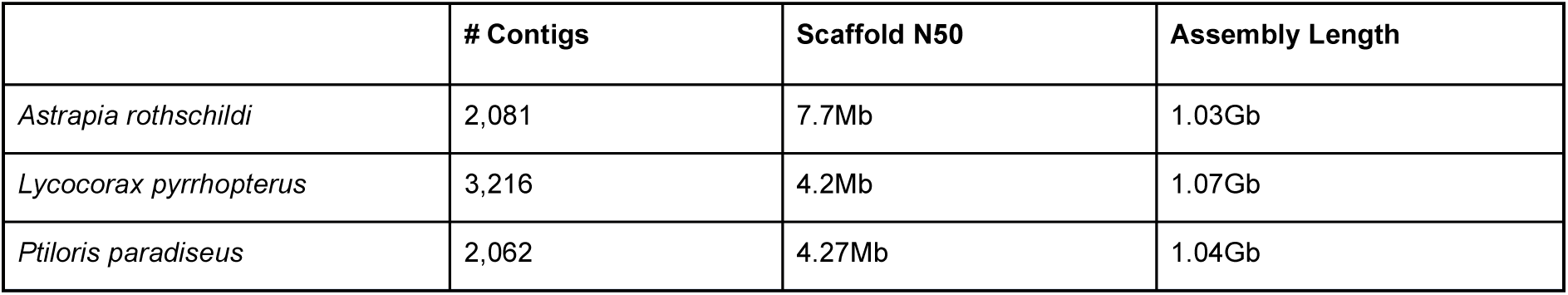
*De novo* Assembly Statistics.

|  | # Contigs | Scaffold N50 | Assembly Length |
| --- | --- | --- | --- |
| *Astrapia rothschildi* | 2,081 | 7.7Mb | 1.03Gb |
| *Lycocorax pyrrhopterus* | 3,216 | 4.2Mb | 1.07Gb |
| *Ptiloris paradiseus* | 2,062 | 4.27Mb | 1.04Gb |

**Supplementary Table S2:**
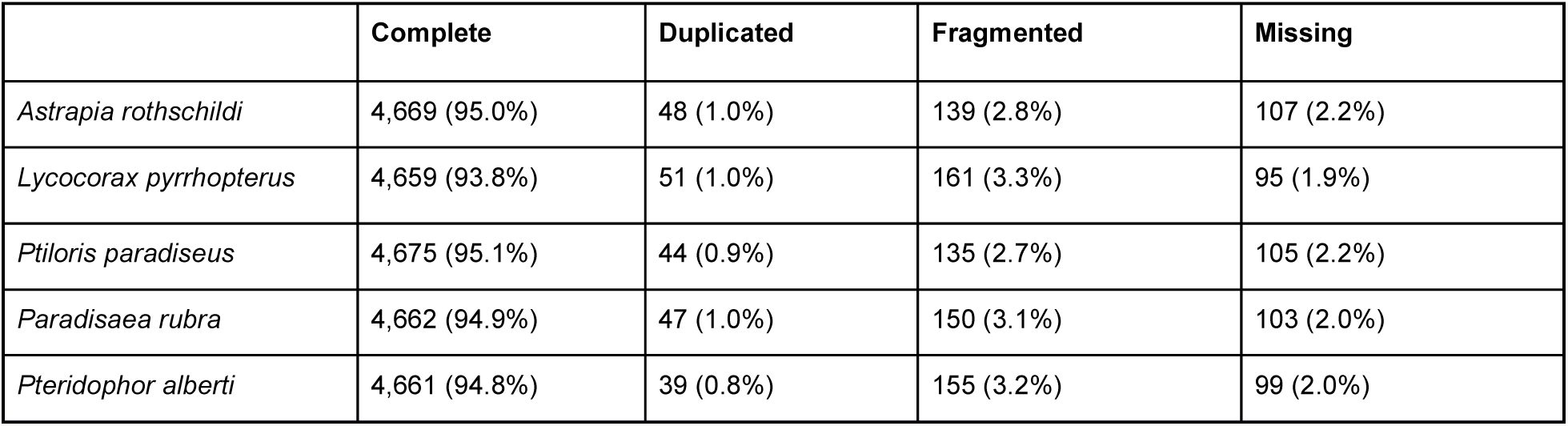
BUSCO scores. Scores were calculated using Busco2 and the aves_odb9 data set (4,915 genes total).

|  | Complete | Duplicated | Fragmented | Missing |
| --- | --- | --- | --- | --- |
| *Astrapia rothschildi* | 4,669 (95.0%) | 48 (1.0%) | 139 (2.8%) | 107 (2.2%) |
| *Lycocorax pyrrhopterus* | 4,659 (93.8%) | 51 (1.0%) | 161 (3.3%) | 95 (1.9%) |
| *Ptiloris paradiseus* | 4,675 (95.1%) | 44 (0.9%) | 135 (2.7%) | 105 (2.2%) |
| *Paradisaea rubra* | 4,662 (94.9%) | 47 (1.0%) | 150 (3.1%) | 103 (2.0%) |
| *Pteridophor alberti* | 4,661 (94.8%) | 39 (0.8%) | 155 (3.2%) | 99 (2.0%) |

**Supplementary Table S3:** Annotation.

|  | # of Transcripts | Average transcript size (bp) | Average introns size (kb) | Average # of introns per gene |
| --- | --- | --- | --- | --- |
| *Astrapia rothschildi* | 16,260 | 1,603 | 2.2 | 9 |
| *Lycocorax pyrrhopterus* | 17,023 | 1,572 | 2.2 | 9 |
| *Ptiloris paradiseus* | 17,269 | 1,584 | 2.2 | 9 |
| *Paradisaea rubra* | 16,822 | 1,561 | 2.2 | 9 |
| *Pteridophor alberti* | 16,721 | 1,562 | 2.2 | 9 |

**Supplementary Table S4:** RepeatMasker annotation of the three birds-of-paradise genome assemblies using a library of our *de novo* repeat annotations of birds-of-paradise merged with existing avian repeat libraries.

|  | *Astrapia rothschildi* |  |  | *Lycocorax pyrrhopterus* |  |  | *Ptiloris paradisaeus* |  |  |
| --- | --- | --- | --- | --- | --- | --- | --- | --- | --- |
| Repeat type | Copies | Total bp | Total % | Copies | Total bp | Total % | Copies | Total bp | Total % |
| **SINE** | 8,019 | 966,992 | 0.09 | 7,974 | 955,190 | 0.09 | 7,977 | 961,945 | 0.09 |
| **LINE** | 128,473 | 38,885,201 | 3.67 | 130,706 | 40,271,136 | 3.76 | 129,094 | 38,994,767 | 3.68 |
| **LTR** | 38,693 | 20,692,445 | 1.95 | 48,395 | 27,819,221 | 2.60 | 39,123 | 21,311,765 | 2.01 |
| **DNA** | 4,582 | 790,017 | 0.07 | 4,734 | 845,120 | 0.08 | 4,617 | 790,421 | 0.07 |
| **Unclassified** | 34,049 | 9,005,494 | 0.85 | 37,519 | 9,167,482 | 0.86 | 30,814 | 8,931,323 | 0.84 |
| **Total inter-spersed repeats** | **213,816** | **70,340,149** | **6.63** | **229,328** | **79,058,149** | **7.39** | **211,625** | **70,990,221** | **6.69** |
| **Small RNA** | 538 | 46,523 | 0.00 | 577 | 50,738 | 0.00 | 546 | 47,744 | 0.00 |
| **Satellites** | 2,884 | 623,756 | 0.06 | 2,706 | 572,161 | 0.05 | 2,855 | 646,354 | 0.06 |
| **Simple repeats** | 195,600 | 9,348,199 | 0.88 | 193,765 | 9,101,884 | 0.85 | 197,648 | 9,318,098 | 0.88 |
| **Low complexity** | 43,076 | 2,388,544 | 0.23 | 42,067 | 2,292,784 | 0.21 | 42,546 | 2,360,570 | 0.22 |
| **Total tandem repeats** | 242,098 | 12,407,022 | 1.17 | 239,115 | 12,017,567 | 1.11 | 243,595 | 12,372,766 | 1.16 |
| **Total repeats** | 455,914 | 82,747,171 | 7.80 | 468,443 | 91,075,716 | 8.50 | 455,220 | 83,362,987 | 7.85 |
| **Assembly** |  | 1.06Gb |  |  | 1.07Gb |  |  | 1.06Gb |  |
| **Gap ('N') bp** |  | 13,196,877 |  |  | 10,466,138 |  |  | 10,993,394 |  |

**Supplementary Table S5:** Characteristics of the manually curated TE consensus sequences from *Lycocorax pyrrhopterus*, including lineage-specific LTR families termed as ‘lycPyrLTR*’.

| Class | Sub-class | Super-family | Family | Subfamily | Similarity to known repeats | Consensus status | Consensus length | TSD |
| --- | --- | --- | --- | --- | --- | --- | --- | --- |
| Retrotransposon | LTR | ERV1 | lycPyrLTR1 | lycPyrLTR1 | None | Complete | 463 | 4 bp |
| Retrotransposon | LTR | ERV1 | lycPyrLTR2 | lycPyrLTR2 | None | Complete | 535 | 4 bp |
| Retrotransposon | LTR | ERV1 | lycPyrLTR3 | lycPyrLTR3 | None | Complete | 623 | 4 bp |
| Retrotransposon | LTR | ERV1 | TguERV3 | TguERV3_LTR2b-L_lycPyr | Partially TguERV3_LTR 2b (68% similarity) | Complete | 601 | 4 bp |
| Retrotransposon | LTR | ERV1 | TguERV1 | TguERV1_LTR1a-L_lycPyr | Partially TguERV1_LTR 1a (72% similarity) | Complete | 600 | 4 bp |
| Retrotransposon | LTR | ERV1 | TguLTR11 | TguLTR11I-L_lycPyr.inc | Partially TguLTR11I + TguERV2_I + TguERV1_I + TguERV3_I (79% + 65% + 63% + 66% similarity) | Incomplete 3' end | 4535 | ? |
| Retrotransposon | LTR | ERV1 | TguLTR12 | TguLTR12-L_lycPyr.inc | Partially TguLTR12 (84% similarity) | Incomplete TSD | 625 | ? |
| Retrotransposon | LTR | ERV2 | lycPyrLTRK1 | lycPyrLTRK1a | None | Complete | 366 | 6 bp |
| Retrotransposon | LTR | ERV2 | lycPyrLTRK1 | lycPyrLTRK1b | None | Complete | 366 | 6 bp |
| Retrotransposon | LTR | ERV2 | lycPyrLTRK2 | lycPyrLTRK2 | None | Complete | 647 | 6 bp |
| Retrotransposon | LTR | ERV2 | lycPyrLTRK3 | lycPyrLTRK3a | None | Complete | 689 | 6 bp |
| Retrotransposon | LTR | ERV2 | lycPyrLTRK3 | lycPyrLTRK3b | None | Complete | 744 | 6 bp |
| Retrotransposon | LTR | ERV2 | lycPyrLTRK4 | lycPyrLTRK4 | None | Complete | 605 | 6 bp |
| Retrotransposon | LTR | ERV2 | lycPyrLTRK5 | lycPyrLTRK5 | None | Complete | 397 | 6 bp |
| Retrotransposon | LTR | ERV2 | lycPyrLTRK6 | lycPyrLTRK6 | None | Complete | 666 | 6 bp |
| Retrotransposon | LTR | ERV2 | lycPyrLTRK7 | lycPyrLTRK7 | None | Complete | 334 | 6 bp |
| Retrotransposon | LTR | ERV2 | lycPyrLTRK8 | lycPyrLTRK8 | None | Complete | 408 | 6 bp |
| Retrotransposon | LTR | ERV2 | lycPyrLTRK9 | lycPyrLTRK9_LTR | None | Complete | 380 | 6 bp |
| Retrotransposon | LTR | ERV2 | lycPyrLTRK9 | lycPyrLTRK9_L.inc | None | Incomplete 3' end | 537 | 6 bp |
| Retrotransposon | LTR | ERV3 | lycPyrLTRL1 | lycPyrLTRL1 | None | Complete | 1171 | 5 bp |
| Retrotransposon | LTR | ERV3 | lycPyrLTRL2 | lycPyrLTRL2 | None | Complete | 1105 | 5 bp |
| Retrotransposon | LTR | ERV3 | lycPyrLTRL3 | lycPyrLTRL3 | None | Complete | 460 | 5 bp |
| Retrotransposon | LTR | ERV3 | lycPyrLTRL4 | lycPyrLTRL4 | None | Complete | 807 | 5 bp |
| Retrotransposon | LTR | ERV3 | lycPyrLTRL5 | lycPyrLTRL5 | None | Complete | 1281 | 5 bp |
| Retrotransposon | LTR | ERV3 | lycPyrLTRL6 | lycPyrLTRL6 | None | Complete | 670 | 5 bp |
| Retrotransposon | LTR | ERV3 | lycPyrLTRL7 | lycPyrLTRL7.inc | Partially Tgu_rep3 (80% similarity) | Incomplete 3' end | 177 | ? |
| Retrotransposon | LTR | ERV3 | TguERVL2 | TguERVL2b-LTR-L_ycPyr | Partially TguERVL2b3_LTR + TguERVL2b1_LTR (85% + 79% similarity) | Complete | 579 | 5 bp |
| Retrotransposon | LTR | ERV3 | TguERVL2 | TguERVL2a2-LTR-L_ycPyr | Partially TguERVL2a2-LTR (93% similarity) | Complete | 941 | 5 bp |
| Retrotransposon | LTR | ERV3 | TguLTRL1 | TguLTRL1-La_ycPyr | Partially TguLTRL1a7 (75% similarity) | Complete | 647 | 5 bp |
| Retrotransposon | LTR | ERV3 | TguLTRL1 | TguLTRL1-Lb_lycPyr.inc | Partially TguERV1_I + TguLTRL1a6 + TguLTRL1a7 (88% + 77% + 74% similarity) | Incomplete 5' end | 2729 | ? |
| Retrotransposon | LTR | ERV3 | TguLTRL1 | TguLTRL1-Lc_lycPyr.inc | Partially TguERV1_I + TguERV2_I + TguLTRL6b (80% + 78% + 97% similarity) | Incomplete 5' and 3' ends | 1754 | ? |
| Retrotransposon | LTR | ERV3 | TguLTRL1 | TguLTRL1-Ld_lycPyr.inc | Partially TguLTRL1a6 + TguLTRL1a7 + TguLTRL1_I (77% + 75% + 75% similarity) | Incomplete 3' end | 2655 | ? |
| Retrotransposon | LTR | ERV3 | TguLTRL1 | TguLTRL1-Le_lycPyr.inc | Partially TguLTRL1a7 + TguERV1_I (75% + 77% similarity) | Incomplete 5' and 3' ends | 3079 | ? |
| Unknown | Unknown | Unknown | Unknown | lycPyr5-275.3inc | None | Incomplete 3' end | 177 | ? |
| Unknown | Unknown | Unknown | Unknown | lycPyr5-1942.inc | None | Incomplete 5' and 3' ends | 620 | ? |
| Unknown | Unknown | Unknown | Unknown | lycPyr6-947.inc_sat | None | Incomplete 5' and 3' ends | 3602 | ? |

**Supplementary Table S6:** Top 10 gene tree topology counts (423 total topologies in 4,450 rooted gene trees). Average Robinson-Foulds distance for all 4,656 gene trees is 3.92. Z: Zebra finch; F: Collared Flycatcher; C: Crow; L: Lycocorax; Pte: Pteridophor; Pti: Ptiloris; Par: Paradisaea; A: Astrapia.

| Topology | Count |
| --- | --- |
| ((Z,F),(C,(L,(Pte,(Pti,(Par,A)))))) | 430 |
| ((Z,F),(C,(L,(Pte,((Pti,Par),A)))) | 357 |
| ((Z,F),(C,(L,(Pte,(Par,(Pti,A)))))) | 279 |
| ((Z,F),(C,(L,(Pti,(Pte,(Par,A)))))) | 224 |
| ((Z,F),(C,(L,((Pti,Par),(Pte,A)))))) | 167 |
| ((Z,F),(C,(L,(Pti,((Pte,Par),A)))) | 166 |
| ((Z,F),(C,(L,(((Pti,Par),Pte),A)))) | 162 |
| ((Z,F),(C,(L,((Pte,Pti),(Par,A)))))) | 161 |
| ((Z,F),(C,(L,((Pti,(Pte,Par)),A)))) | 159 |
| ((Z,F),(C,(L,(Pti,(Par,(Pte,A)))))) | 156 |

**Supplementary Table S7:** Saturation Analysis. Pairwise dn/ds ratio.

|  |  |  |  |  |  |  |  |
| --- | --- | --- | --- | --- | --- | --- | --- |
| **Astrapia** |  |  |  |  |  |  |  |
| **Bilscor** | 0.036 |  |  |  |  |  |  |
| **Ensfalt** | 0.046 | 0.063 |  |  |  |  |  |
| **Enstgut** | 0.044 | 0.059 | 0.046 |  |  |  |  |
| **Lycocorax** | 0.014 | 0.037 | 0.046 | 0.044 |  |  |  |
| **Paradisaea** | 0.006 | 0.034 | 0.046 | 0.043 | 0.014 |  |  |
| **Pteridophor** | 0.007 | 0.036 | 0.046 | 0.044 | 0.014 | 0.007 |  |
| **Ptiloris** | 0.006 | 0.036 | 0.046 | 0.044 | 0.014 | 0.006 | 0.007 |

**Supplementary Table S8.** Genes under positive selection.

| GenBank Accession | Gene Symbol |
| --- | --- |
| XM_016302299 | RSPH14 |
| XM_005060676 | SNX18 |
| XM_005061576 | RSG1 |
| XM_016299097 | COL4A1 |
| XM_016304880 | MCTP1 |
| XM_005042552 | NDRG1 |
| XM_005062657 | MRPL34 |
| XM_005057187 | BPIFB2 |
| XM_016298377 | LOC101809528 |
| XM_005057074 | LOC101821424 |
| XM_016301269 | LOC101807976 |
| XM_016302077 | TRAFD1 |
| XM_005048141 | *SH2D4B* |
| XM_005044853 | MGARP |
| XM_005039384 | ADAMTS20 |
| XM_005037072 | TAF10 |
| XM_005054309 | C14H16orf71 |
| XM_005056583 | EVI2A |
| XM_005038551 | CCDC181 |
| XM_016298245 | COL4A5 |
| XM_005059334 | LAD1 |
| XM_016301828 | LOC101813372 |
| XM_016296212 | AGAP3 |
| XM_005051230 | FETUB |
| XM_005043410 | POLH |
| XM_005054364 | DRC3 |
| XM_016300815 | ZWILCH |
| XM_005043438 | WDR27 |
| XM_005058894 | S100A11 |
| XM_005046922 | C5H11orf74 |
| XM_016306090 | SLC9A2 |
| XM_005050076 | LOC101808676 |
| XM_016300582 | ATP7B |
| XM_005051988 | TCF12 |
| XM_016303963 | LOC107604184 |
| XM_005061593 | LOC101821569 |
| XM_016298934 | ITPK1 |
| XM_016302041 | CARHSP1 |
| XM_005049042 | LOC101807582 |
| XM_016303572 | IDO2 |
| XM_005052727 | LOC101813437 |
| XM_005047366 | GPATCH2L |
| XM_005057073 | WISP2 |
| XM_005055863 | PMP22 |
| XM_005062304 | IGSF21 |
| XM_005050061 | SLC5A9 |
| XM_016298968 | CABP4 |
| XM_005056673 | LOC101816855 |
| XM_005056783 | LOC101817428 |
| XM_005056335 | CBX2 |
| XM_005042325 | PTDSS1 |
| XM_005055795 | BARHL1 |
| XM_005048773 | HABP2 |
| XM_016303687 | SYTL1 |
| XM_016303706 | PAQR7 |
| XM_016301660 | CCDC89 |
| XM_016305278 | KIF3C |
| XM_005059500 | PHLDA3 |
| XM_005039461 | PHLDA1 |
| XM_016299105 | GPX2 |
| XM_005046697 | GPX2 |
| XM_005049721 | DTX3L |
| XM_005039251 | LOC101813208 |
| XM_005049817 | RNASEL |
| XM_005050542 | GPR88 |
| XM_005045976 | NEXMIF |
| XM_005061804 | LOC101813871 |
| XM_005050079 | MOB3C |
| XM_005043012 | ITPKB |
| XM_005061100 | LOC101811548 |
| XM_005052526 | SLC12A3 |
| XM_005060823 | LOC101822159 |
| XM_005060810 | DOCK8 |
| XM_016298010 | *SLC7A2* |
| XM_016299897 | G6PC2 |
| XM_005040638 | *ZEB1* |
| XM_005045593 | CCDC149 |
| XM_005055395 | *ALAD* |
| XM_005048989 | *SPAG16* |
| XM_005060340 | HAUS1 |
| XM_005044316 | SLC25A27 |
| XM_005055265 | PTPN11 |
| XM_005056255 | SLC25A10 |
| XM_016298524 | LOC101816285 |
| XM_005061480 | TSPAN9 |
| XM_005061498 | TCAF2 |
| XM_016299390 | ANXA11 |
| XM_005059540 | TAF11 |
| XM_016304488 | NCLN |
| XM_005051635 | NR2E3 |
| XM_005059699 | P3H4 |
| XM_005038313 | GAB2 |
| XM_016300131 | AGL |
| XM_016296086 | TERT |
| XM_016303624 | FGFR1 |
| XM_005055228 | GCN1 |
| XM_005037507 | LIMS1 |
| XM_005058597 | PAFAH1B2 |
| XM_005060625 | LOC101821368 |
| XM_005059102 | UBE2Q1 |
| XM_016297193 | MZT1 |
| XM_005038138 | B3GLCT |
| XM_005038022 | PDS5B |
| XM_016300038 | LMO4 |
| XM_005050724 | RBP2 |
| XM_005053530 | CYFIP2 |
| XM_005039572 | STRA8 |
| XM_005042040 | *TAF4B* |
| XM_005042005 | TUBB6 |
| XM_005048677 | SH3PXD2A |
| XM_005048419 | *IDE* |
| XM_005047986 | DNAJB12 |
| XM_005038401 | GSTK1 |
| XM_005046991 | DNAL1 |
| XM_005049866 | *MYOC* |
| XM_005050007 | PLPP6 |
| XM_005053617 | NHP2 |
| XM_005042611 | LOC101815973 |
| XM_005051099 | OSTN |
| XM_005051315 | *MECOM* |
| XM_005037953 | CDADC1 |
| XM_016303810 | RPS25 |
| XM_016299107 | TCIRG1 |
| XM_005039906 | ACSS3 |
| XM_016304331 | MYF5 |
| XM_005045389 | ARAP2 |
| XM_005045346 | UBE2K |
| XM_005054795 | SPECC1L |
| XM_005054910 | MAPK1 |
| XM_005043131 | LOC101809314 |
| XM_016297079 | IMPG1 |
| XM_005054684 | TSR3 |
| XM_005043198 | LOC101822112 |
| XM_005043614 | SLC22A2 |
| XM_005054008 | RAB26 |
| XM_005037142 | SH3BGR |
| XM_016298043 | RUNX1 |
| XM_005037183 | *RIPK4* |
| XM_005045614 | LOC101812359 |
| XM_005061387 | NACA |
| XM_005046816 | *CTSD* |
| XM_005047384 | ERH |
| XM_005040064 | LOC101810161 |
| XM_005048086 | *PAPSS2* |
| XM_016301575 | ATP10B |
| XM_016300850 | *SPATA5L1* |
| XM_016300780 | AKAP13 |
| XM_005039138 | CBLL1 |
| XM_016303057 | RELN |
| XM_016304620 | GNG10 |
| XM_005057597 | GNB1 |
| XM_005038109 | LOC101817904 |
| XM_005053334 | TRH |
| XM_005037520 | LOC101809883 |
| XM_005048986 | ATIC |
| XM_005059186 | KCND3 |
| XM_005059571 | LOC101810386 |
| XM_016298945 | LOC107603674 |
| XM_016298921 | LOC101813292 |
| XM_005057359 | TAF4 |
| XM_005046501 | PKP3 |
| XM_005056669 | ALDH3A2 |
| XM_005039746 | BAIAP2L2 |
| XM_005049157 | WDR12 |
| XM_016298059 | RFC1 |
| XM_016296127 | *MTBP* |
| XM_005045169 | SPCS3 |
| XM_005059547 | *RPRML* |
| XM_005048151 | RASGEF1A |
| XM_016305207 | NTRK2 |
| XM_005059489 | *APOBEC2* |
| XM_016304871 | ZCCHC7 |
| XM_016304884 | CENPK |
| XM_016302782 | *DNAH9* |
| XM_005059010 | *DCST2* |

**Supplementary Table S9.**
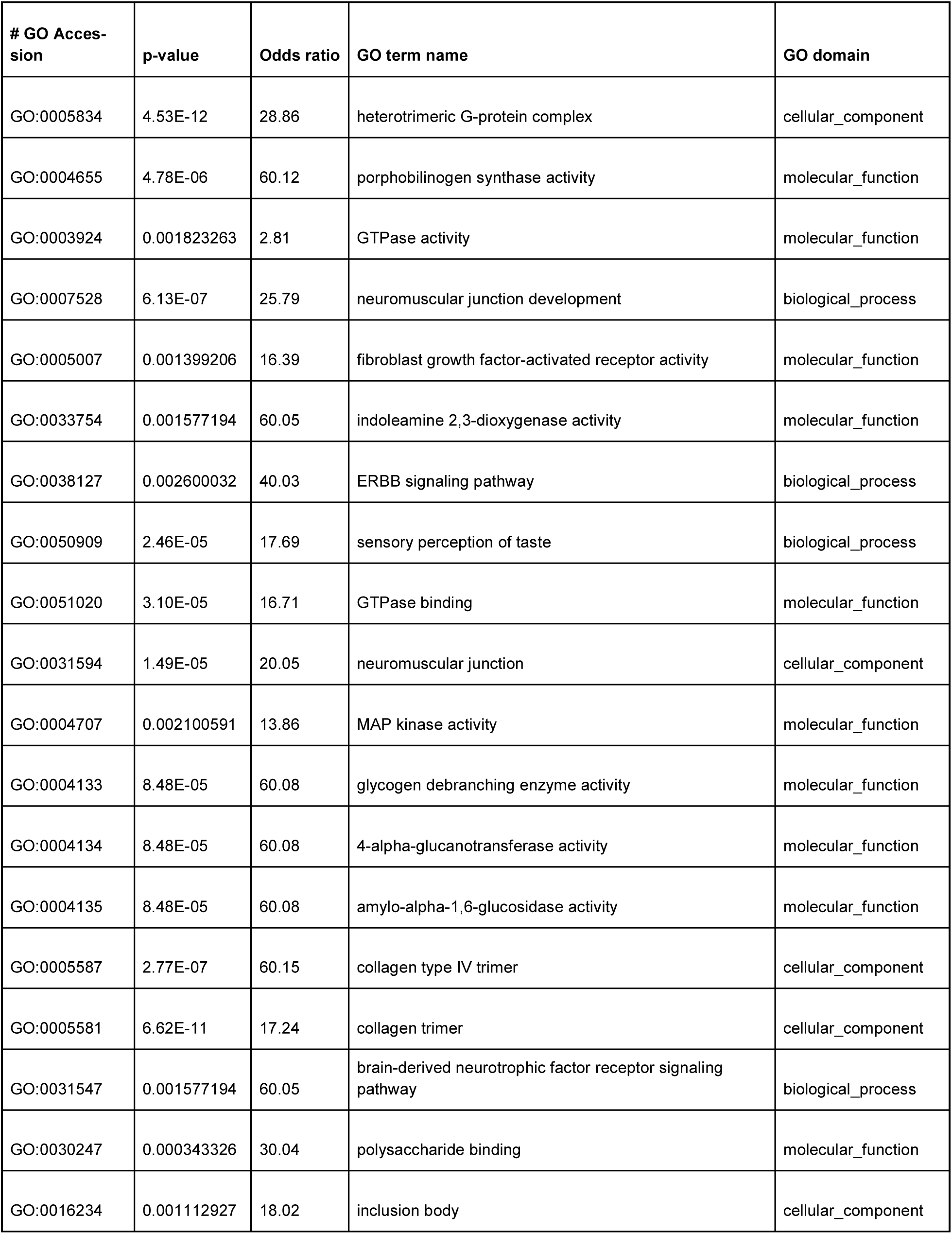
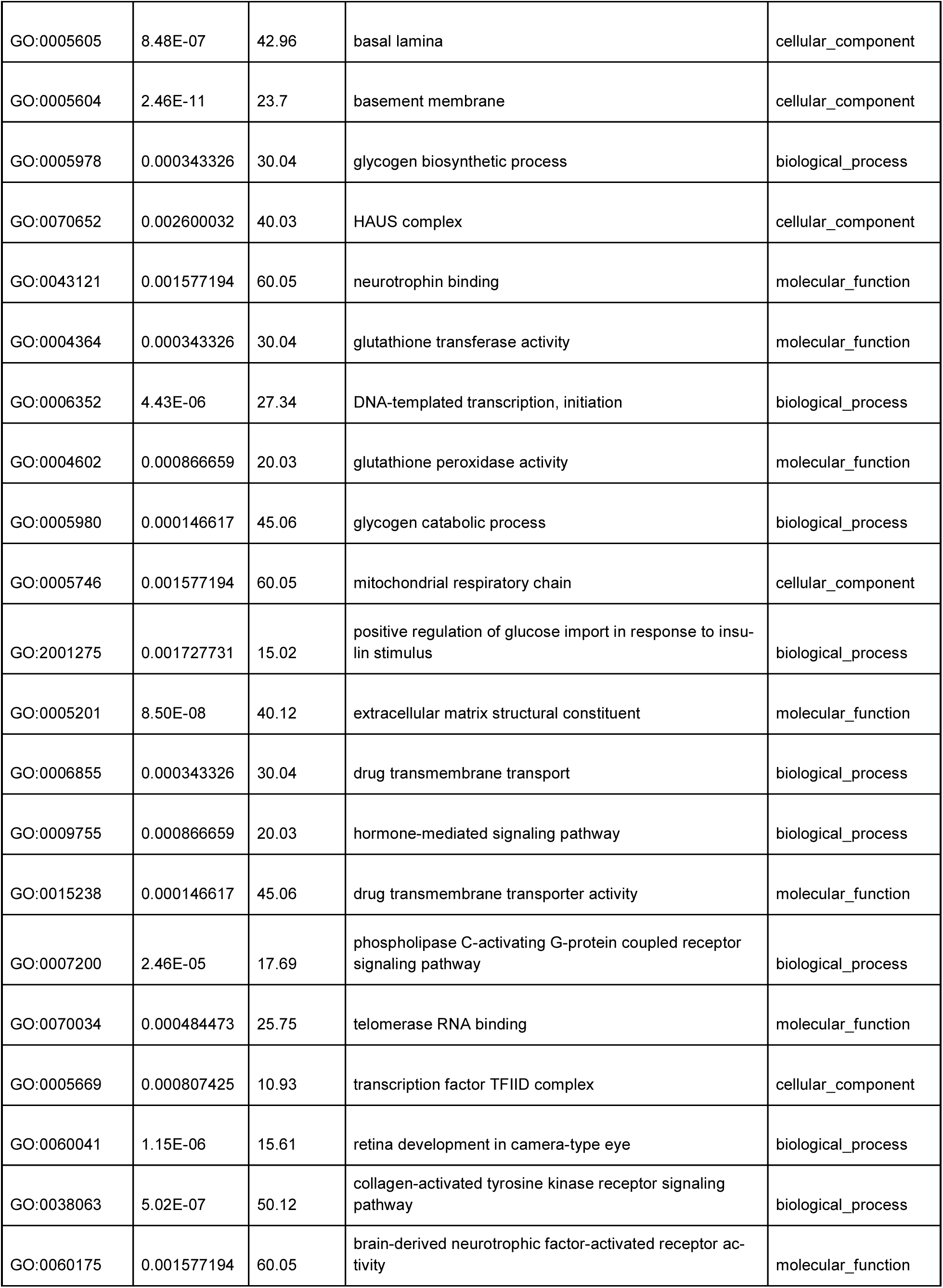
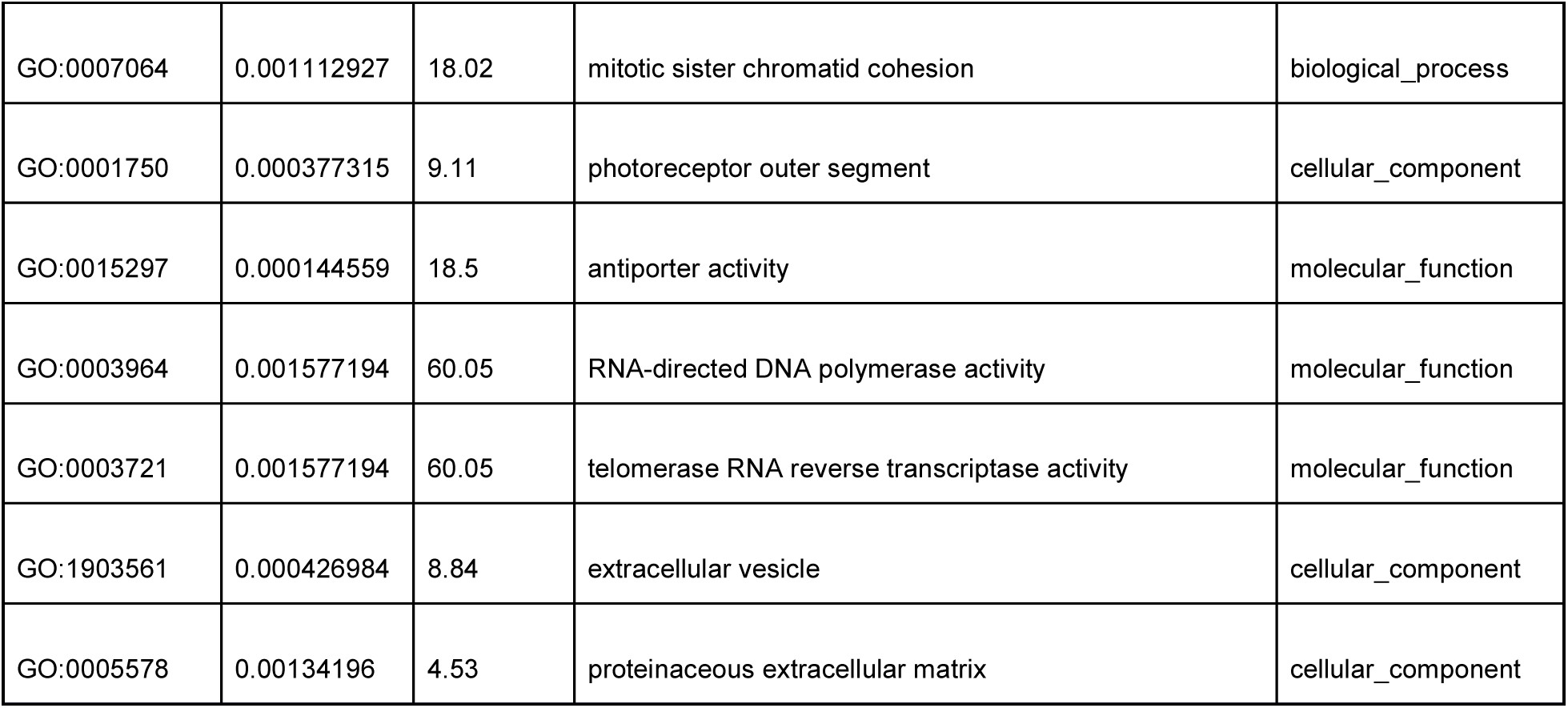
Enriched GO terms in the positive selection analysis.

**Supplementary Table S10:** Summary of gene gain and loss events inferred after correcting for annotation and assembly error across all 13 species. The number of rapidly evolving families is shown in parentheses for each type of change.

|  | Expansions |  |  | Contractions |  |  | No Change | Avg. Expansion |
| --- | --- | --- | --- | --- | --- | --- | --- | --- |
|  | Families | Genes gained | genes/ expansion | Families | Genes lost | genes/ contraction |  |  |
| Paradisaea | 248 (40) | 297 | 1.2 | 209 (3) | 215 | 1.03 | 8555 | 0.009323 |
| Astrapia | 314 (40) | 398 | 1.27 | 455 (31) | 543 | 1.19 | 8243 | -0.016537 |
| Ficedula | 329 (23) | 480 | 1.46 | 560 (7) | 671 | 1.2 | 8123 | -0.020977 |
| Lycocorax | 513 (16) | 612 | 1.19 | 338 (2) | 358 | 1.06 | 8161 | 0.027747 |
| Taeniopygia | 1463 (17) | 2009 | 1.37 | 977 (7) | 1040 | 1.06 | 6572 | 0.091565 |
| Ptiloris | 334 (49) | 401 | 1.2 | 203 (5) | 219 | 1.08 | 8475 | 0.020200 |
| Pteridophor | 241 (13) | 274 | 1.14 | 297 (6) | 309 | 1.04 | 8474 | -0.002997 |
| Crow | 362 (6) | 480 | 1.33 | 1708 (45) | 2050 | 1.2 | 6942 | -0.172475 |

**Supplementary Table S11:** Assembly/Annotation error estimation and gene gain/loss rates in a single *λ* model in the 13 mammals included in this study compared to previous studies using fewer species.

| | $\lambda$ (No Error Model) | $\varepsilon$ (Estimated error) | $\lambda$ (Error Model = $\varepsilon$) |
| --- | --- | --- | --- |
| **8 bird species in this study** | 0.00221 | 0.01025 | 0.00205 |
| **12 *Drosophila* species\*** | 0.00121 | 0.04102 | 0.00059 |
| **10 mammal species\*** | 0.00238 | 0.07324 | 0.00186 |
| **16 fungi species\*** | 0.0008 | 0.02771 | 0.00061 |

\* Dataset from Han et al. 2013 [1].

**Supplementary Table S12:** Enriched GO terms in rapidly evolving birds-of-paradise families. The number in parentheses for Rapidly evolving lineages indicates the extent of change along that lineage (ie Astrapia (+6) means that the Astrapia lineage gained 6 genes). Lineages within the BOP clade are indicated by bold text. See Figure S1 for internal node labels.

| Family ID | GO accession: GO name | Rapidly evolving lineages | Enriched after FDR correction? |
| --- | --- | --- | --- |
| 1 | GO:0001077: transcriptional activator activity, RNA polymerase II core promoter proximal region sequence-specific binding | **Astrapia (+6)**<br>BOP13 (-17)<br>BOP1 (+9)<br>Taeniopygia (+36)<br>**Pteridophor (-4)**<br>Crow (-12) | * |
| 1 | GO:0001078: transcriptional repressor activity, RNA polymerase II core promoter proximal region sequence-specific binding | **Astrapia (+6)**<br>BOP13 (-17)<br>BOP1 (+9)<br>Taeniopygia (+36)<br>**Pteridophor (-4)**<br>Crow (-12) |  |
| 1 | GO:0000978: RNA polymerase II core promoter proximal region sequence-specific DNA binding | **Astrapia (+6)**<br>BOP13 (-17)<br>BOP1 (+9)<br>Taeniopygia (+36)<br>**Pteridophor (-4)**<br>Crow (-12) | * |
| 1 | GO:0000977: RNA polymerase II regulatory region sequence-specific DNA binding | **Astrapia (+6)**<br>BOP13 (-17)<br>BOP1 (+9)<br>Taeniopygia (+36)<br>**Pteridophor (-4)**<br>Crow (-12) | * |
| 1 | GO:0001223: transcription coactivator binding | **Astrapia (+6)**<br>BOP13 (-17)<br>BOP1 (+9)<br>Taeniopygia (+36)<br>**Pteridophor (-4)**<br>Crow (-12) |  |
| 13 | GO:0004984: olfactory receptor activity | **Astrapia (+6)**<br>**Lycocorax (-6)**<br>Taeniopygia (+17)<br>**BOP9 (+5)**<br>Crow (-9) | * |
| 26 | GO:0005200: structural constituent of cytoskeleton | **BOP11 (-26)**<br>Ficedula (+16)<br>BOP13 (-21)<br>BOP1 (+12)<br>Taeniopygia (+20)<br>**BOP9 (-3)** | * |
| 30 | GO:0001948: glycoprotein binding | **Paradisaea (+6)**<br>**Astrapia (-3)** |  |
| 30 | GO:0005041: low-density lipoprotein receptor activity | **Paradisaea (+6)**<br>**Astrapia (-3)** |  |
| 31 | GO:0001964: startle response | **Paradisaea (+3)**<br>**BOP9 (+5)** |  |
| 31 | GO:0008344: adult locomotory behavior | **Paradisaea (+3)**<br>**BOP9 (+5)** | * |
| 36 | GO:0005112: Notch binding | **Ptiloris (+3)** | * |
| 39 | GO:0005198: structural molecule activity | **BOP9 (-3)** | * |
| 49 | GO:0003777: microtubule motor activity | **Astrapia (+3)**<br>Crow (+15) | * |
| 65 | GO:0004222: metalloendopeptidase activity | **Astrapia (+4)**<br>Ptiloris (-3) | * |
| 67 | GO:0005216: ion channel activity | **BOP11 (+4)**<br>Crow (-4) | * |
| 76 | GO:0001657: ureteric bud development | **BOP11 (+3)**<br>Crow (-4) | * |
| 97 | GO:0003958: NADPH-hemoprotein reductase activity | **Astrapia (+3)** | * |
| 97 | GO:0004517: nitric-oxide synthase activity | **Astrapia (+3)** | * |
| 97 | GO:0005272: sodium channel activity | **Astrapia (+3)** | * |
| 102 | GO:0007155: cell adhesion | **Ptiloris (-2)**<br>Crow (+6) | * |
| 121 | GO:0060348: bone development | **Astrapia (-4)** |  |
| 121 | GO:0002091: negative regulation of receptor internalization | **Astrapia (-4)** | * |
| 130 | GO:0005201: extracellular matrix structural constituent | **Paradisaea (+3)** |  |
| 138 | GO:0014719: skeletal muscle satellite cell activation | **BOP5 (+1)** | * |
| 170 | GO:0006955: immune response | **Astrapia (-2)**<br>Taeniopygia (+8)<br>**Ptiloris (+2)**<br>**BOP5 (-3)**<br>Crow (-4) | * |
| 252 | GO:0019992: diacylglycerol binding | **BOP11 (+2)**<br>**Pteridophor (+2)** | * |
| 290 | GO:0030234: enzyme regulator activity | **Paradisaea (-3)** |  |
| 312 | GO:0003956: NAD(P)+-protein-arginine ADP-ribosyltransferase activity | **Paradisaea (+2)**<br>Ficedula (+5)<br>Taeniopygia (-3)<br>**Ptiloris (+2)**<br>**BOP7 (+1)**<br>Crow (-3) |  |
| 321 | GO:0017112: Rab guanyl-nucleotide exchange factor activity | **Ptiloris (-2)**<br>**Pteridophor (+2)** | * |
| 477 | GO:0050699: WW domain binding | **BOP11 (+2)** |  |
| 584 | GO:0004415: hyaluronoglucosaminidase activity | **Lycocorax (+4)** | * |
| 637 | GO:0005149: interleukin-1 receptor binding | **Astrapia (+2)**<br>**BOP9 (-2)** | * |
| 836 | GO:0017080: sodium channel regulator activity | **Paradisaea (+3)**<br>**BOP9 (-2)** |  |
| 867 | GO:0002060: purine nucleobase binding | **Paradisaea (+2)**<br>**Ptiloris (+2)** | * |
| 867 | GO:0004645: phosphorylase activity | **Paradisaea (+2)**<br>**Ptiloris (+2)** | * |

